# Arf GTPase activates the WAVE Regulatory Complex through a novel binding site

**DOI:** 10.1101/2022.05.13.491857

**Authors:** Sheng Yang, Yijun Liu, Abbigale Brown, Matthias Schaks, Bojian Ding, Daniel A. Kramer, Li Ding, Olga Alekhina, Daniel D. Billadeau, Saikat Chowdhury, Junmei Wang, Klemens Rottner, Baoyu Chen

## Abstract

Crosstalk between Rho- and Arf-family GTPases plays an important role in linking actin cytoskeletal remodeling to membrane protrusion, organelle structure, and vesicle trafficking. The central actin regulator, WAVE Regulatory Complex (WRC), is a converging point of Rac1 (a Rho-family GTPase) and Arf signaling in many processes, but how Arf promotes WRC activation is unknown. Here we reconstituted a direct interaction between Arf and WRC. This interaction can be greatly enhanced by Rac1 binding to the D site of the WRC. Arf1 binds to a newly identified conserved surface on Sra1 located between the D site and the WH2 helix of WAVE1, which can drive WRC activation using a mechanism distinct from that of Rac1. Mutating Arf binding site abolishes Arf1-WRC interaction, disrupts Arf1-mediated WRC activation, and impairs lamellipodia morphology. This work uncovers a new mechanism underlying WRC activation and provides a mechanistic foundation for studying how WRC-mediated actin polymerization links Arf and Rac signaling in the cell.

## Introduction

Small GTPases of the Ras superfamily control diverse processes throughout eukaryotic cells (Wennerberg et al., 2005). Among them, the distantly related Arf-family and Rho-family GTPases play distinct roles and yet have extensive crosstalk in many different processes. Arf GTPases are key players in various steps of membrane trafficking and organelle morphogenesis, where they are best known to promote the assembly of coat proteins to initiate vesicle formation (D’Souza-Schorey and Chavrier, 2006; Donaldson and Jackson, 2011; Gillingham and Munro, 2007; Sztul et al., 2019). Rho GTPases, such as Rac1, are central regulators of the actin cytoskeleton in the formation of various cell membrane protrusions, such as lamellipodia and filopodia, where they are best known to promote cell migration, adhesion, and endocytosis (Etienne-Manneville and Hall, 2002; Mosaddeghzadeh and Ahmadian, 2021). Since it was discovered over two decades ago (Boshans et al., 2000; D’Souza-Schorey et al., 1997; Radhakrishna et al., 1999; Santy and Casanova, 2001), the crosstalk between Arf- and Rac1-mediated signaling pathways has been recognized as an essential component for the regulation of actin cytoskeletal dynamics during cell migration, spreading, adhesion, fusion, phagocytosis, and endocytosis (Boshans et al., 2000; Chen et al., 2003; D’Souza-Schorey et al., 1997; Hunt et al., 2022; Myers and Casanova, 2008; Phuyal and Farhan, 2019; Radhakrishna et al., 1999; Santy and Casanova, 2001; Singh et al., 2017). Nevertheless, our knowledge of the underlying molecular mechanism remains fragmental.

In addition to the role of Arf in regulating phospholipid microenvironment (Honda et al., 1999; Krauss et al., 2003), endosomal recycling of Rac1 (Balasubramanian et al., 2007; Boshans et al., 2000; Radhakrishna et al., 1999), and the localization and activity of Rac1 GEFs (Chen et al., 2003; Koo et al., 2007; Palacios et al., 2002; Santy et al., 2005), GAPs(Hu et al., 2009), and adaptors (D’Souza-Schorey et al., 1997; Tarricone et al., 2001), a plethora of studies have underscored the convergence of Arf and Rac1 on a central actin nucleation promotion factor known as the WAVE Regulatory Complex (WRC) in many processes (Humphreys et al., 2012b, 2012a, 2013, 2016; Koronakis et al., 2011; Lewis-Saravalli et al., 2013; Marchesin et al., 2015; Singh et al., 2019, 2020). The WRC is a 400-kDa protein assembly containing five conserved proteins: Sra1 (or Cyfip2), Nap1 (or Hem1), Abi2 (or Abi1, Abi3), HSPC300, and WAVE1 (or WAVE2, WAVE3, members of the Wiskott-Aldrich Syndrome protein family). In the basal state, the WRC keeps WAVE auto-inhibited in the cytosol by sequestering the WCA (WH2-central-acidic) sequence at the C-terminus of WAVE through a collection of interactions with the Sra1 subunit and the “meander” sequence of WAVE (**Figure 1**, cartoon) (Chen et al., 2014a, 2010; Derivery et al., 2009; Eden et al., 2002; Ismail et al., 2009). Various membrane ligands can directly interact with and recruit the WRC to the plasma membrane and simultaneously activate it to release the WCA, which in turn can bind the Arp2/3 complex to polymerize branched actin filaments (Chen et al., 2014b, 2017; Koronakis et al., 2011; Lebensohn and Kirschner, 2009; Padrick et al., 2008; Rottner et al., 2021; Takenawa and Suetsugu, 2007; Zou et al., 2018). Among these ligands, Rac1 is the canonical activator of the WRC (Rottner et al., 2021). It acts by directly binding to two distinct locations on the opposite ends of the Sra1 subunit, which are named A and D sites, respectively. The two sites have ∼40-fold difference in the affinity for Rac1 (Chen et al., 2017, 2010). The recent cryo-EM structures revealed that Rac1 binding to the low affinity site (A site), but not the high-affinity site (D site), drives a conformational change to allosterically destabilize the WCA leading to WRC activation (Ding et al., 2022).

**Figure 1.**
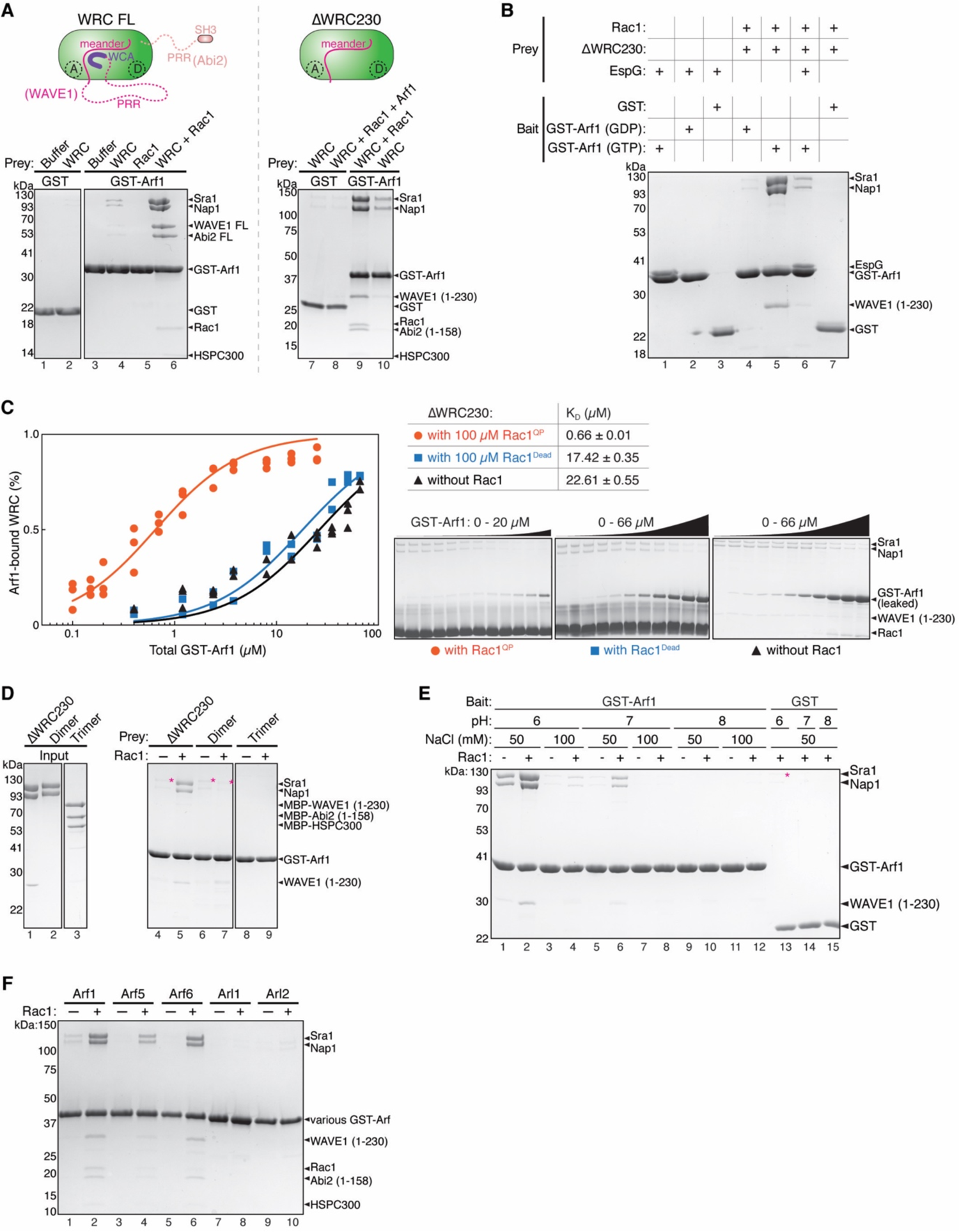
Arf-WRC interaction is direct and can be greatly enhanced by Rac1. **(A)** Coomassie blue-stained SDS-PAGE gels showing GST-Arf1 pull-down of WRC FL (left) and ΔWRC230 (right) in the presence or absence of untagged Rac1^QP^. In the schematic of respective WRCs, dotted lines indicate unstructured sequences. Both the A and D sites for Rac1 binding are indicated. **(B)** Coomassie blue-stained SDS-PAGE gels showing pull-down of ΔWRC230 by GST-Arf1 in indicated nucleotide states or in the presence of the Arf1-binding protein EspG. **(C)** Equilibrium pull-down (EPD) assay to measure the binding affinity of the Arf1-WRC interaction in the presence of indicated Rac1 variants. On the left is the quantification of the data pooled from two to three independent experiments for each condition and globally fitted to obtain the binding isotherms. The derived dissociation constant (K_D_) and fitting errors are shown in the table. On the right are representative Coomassie blue-stained SDS-PAGE gels of the supernatant samples use for quantification. **(D)** Coomassie blue-stained SDS-PAGE gels showing GST-Arf1 pull-down of WRC subcomplexes in the presence or absence of Rac1^QP^. Dimer is the Sra1/Nap1 subcomplex. Trimer is the WAVE1 (1-230)/Abi2(1-158)/HSPC300 subcomplex. Asterisks indicate weak binding signals. **(E)** Arf1 binding to the WRC is sensitive to pH and salt concentration. Shown is Coomassie blue-stained SDS PAGE from GST-Arf1 pull down of ΔWRC230 in indicated buffer conditions, in the presence or absence of Rac1^QP^. Red asterisk indicates increased background binding to GST beads at pH 6, to avoid which we use pH 7 and 50 mM NaCl throughout this study. **(F)** Coomassie blue-stained SDS-PAGE gel showing pull-down of ΔWRC230 by different GST-tagged Arf-family members with or without Rac1^QP^.

The connection between Arf1, Rac1 and the WRC was initially discovered by proteomic and cellular studies to identify proteins important for clathrin-AP-1-coated carrier biogenesis at the trans-Golgi network (TGN) (Anitei et al., 2010; Baust et al., 2006). A more direct connection was established in a seminal study by Koronakis and colleagues in 2011, in which they reconstituted WRC activation using lipid-coated beads and mammalian brain lysates (Koronakis et al., 2011). They found lipid-coated beads containing individual Rac1 or Arf1 only bound and activated WRC weakly, but beads containing both GTPases dramatically enhanced WRC membrane recruitment and activation (Koronakis et al., 2011). After that, a series of studies further corroborated the connection of Arf to the WRC. For example, Arf79 (the Arf1 homolog in *Drosophila*) was found to be critical for Sra1 localization and concomitant formation of lamellipodia (Humphreys et al., 2012a). This function could not be complemented by Rac overexpression, but could be restored by expressing the human Arf1, underlining the importance of Arf1 to WRC activation and the conserved role of the Arf-WRC interaction across species (Humphreys et al., 2012a). Furthermore, two different types of bacterial pathogens, *Salmonella enterica* and enteropathogenic and enterohemorrhagic *Escherichia coli* (EPEC and EHEC), could both hijack the Arf1-Rac1-WRC signaling axis to facilitate infection, albeit with opposite objectives (from the bacteria point of view) and via distinct mechanisms (Humphreys et al., 2012b, 2013, 2016). In addition, the cooperative actions of Arf1 (or Arf6) and Rac1 on the WRC were found to be critical for the migration of invasive MDA-MB-231 breast cancer cells (Lewis-Saravalli et al., 2013; Marchesin et al., 2015). Moreover, a missense mutation in *Hem1* from patients with an inherited immunologic syndrome named immunodeficiency-72 with an autoinflammation (IMD72) was found to disrupt Arf1-, but not Rac1-mediated WRC activation (Cook et al., 2020).

Despite the importance of Arf1-Rac1-WRC signaling in various normal and disease-related processes, the mechanism by which Arf1 achieves this function is unknown. Sharing less than 30% sequence identity with Rac1, Arf1 may use a distinct mechanism to regulate the WRC. But does Arf1 directly interact with the WRC or Rac1 at all? If yes, what is the interaction mechanism, and what is the biochemical and structural basis of the cooperativity between Arf1 and Rac1? To answer these questions, here we have reconstituted a direct interaction between Arf and the WRC in solution by using purified proteins. We find the interaction is greatly enhanced by Rac1 binding to the WRC mainly on the D site. Remarkably, once bound to the WRC, Arf1 can directly activate the WRC independent of Rac1 binding to the A site. We further identified the Arf1 binding site, which is located at a conserved surface on Sra1 between the D site and the W helix of the WCA domain of WAVE. Mutating the Arf1 binding site abolished Arf1 binding, disrupted Arf1-mediated WRC activation, and impaired lamellipodia morphology. Together, our work reveals a new mechanism underlying WRC activation and paves the way for understanding how WRC-mediated actin polymerization integrates signals from Arf and Rac in various processes.

## Results

### Arf GTPases directly interact with WRC, and the interaction is greatly enhanced by Rac1

The interaction between Arf1 and WRC was initially discovered using lipid-coated beads where both Arf1 and Rac1 were anchored on the membrane and incubated with mammalian brain extracts (Koronakis et al., 2011). To examine if the Arf1-WRC interaction is direct and, if yes, to determine the underlying mechanism, we reconstituted this interaction in solution using recombinantly purified proteins. We found that GST-tagged Arf1 could directly pull down both the full-length (FL) WRC and a truncated WRC named ΔWRC230 (**Figure 1A**, lane 4, 10) (Chen et al., 2014a, 2017). ΔWRC230 represents the minimal, structured core of the WRC, since it lacks the C-terminal unstructured proline-rich region (PRR) and the WCA sequence of WAVE1, as well as the unstructured PRR and the SH3 domain of Abi2 (**Figure 1A**, cartoon). Although the binding signals were weak, they were specific in comparison to the background signals in GST controls (**Figure 1A**, lane 2 vs. 4, lane 7 vs. 10). Thus, Arf1 directly interacts with the WRC, and the structured core of the WRC is sufficient to bind Arf1.

To test if and how Rac1 can enhance Arf1 binding to the WRC, we used a Rac1 that contained two mutations, Q61L and P29S, which greatly enhanced Rac1 binding to the WRC as shown in our previous studies (Chen et al., 2017). Unless otherwise noted, we refer to this Rac1^Q61L/P29S^ construct as Rac1 or Rac1^QP^ interchangeably in this study. We found including free Rac1 in the pull-down reactions drastically enhanced GST-Arf1 binding to the WRC (**Figure 1A**, lane 6, 9). Note that Rac1 did not directly interact with Arf1 (**Figure 1A**, lane 5), but was co-retained with WRC by GST-Arf1 (**Figure 1A**, lane 6, 9). These results suggest that Arf1 and Rac1 can simultaneously bind to the same WRC via non-overlapping binding sites, and that Rac1 binding greatly stabilizes Arf1 binding.

As molecular switches, GTPases usually use the GTP state to engage with downstream effector proteins. We found that the Arf1-WRC interaction was also dependent on the nucleotide state of Arf1. Only Arf1 loaded with GTP, but not GDP, showed robust binding (**Figure 1B**, lane 4 vs. 5). Moreover, the interaction could be specifically blocked by EspG, a bacterial effector protein secreted into the host cell by enteropathogenic (EPEC) and enterohaemorrhagic (EHEC) *Escherichia coli* during infection (Dong et al., 2012; Humphreys et al., 2016). EspG directly binds the GTP-form of Arf1 and Arf6 (Dong et al., 2012; Humphreys et al., 2016) (also see **Figure 1B**, lane 1-3). This interaction was suggested to disrupt Arf-WRC signaling in host cells, which helped these extracellular pathogens evade WRC-mediated phagocytosis (Humphreys et al., 2016). Our data suggest EspG can achieve this by directly competing off Arf1 (and/or Arf6) binding to the WRC. Therefore, Arf1 may use the same surface to interact with the WRC and EspG.

We next used our previously established equilibrium pull-down (EPD) assay to quantitatively measure the enhancement of Arf1 binding by Rac1 (Chen et al., 2017; Pollard, 2010) (**Figure 1C**). We found that in the absence of Rac1, the Arf1-WRC interaction was weak, with a dissociate constant K_D_ ∼23 µM (**Figure 1C**, black). By contrast, in the presence of 100 µM Rac1, which should saturate both A and D sites of the WRC (Chen et al., 2017; Ding et al., 2022), Arf1 binding affinity was increased nearly 30 fold (K_D_ ∼ 0.66 µM, **Figure 1C**, orange). The enhanced binding was not an artifact of high concentration of free Rac1 included in the assay, as a mutant Rac1, in which the entire Switch I motif critical for WRC binding was removed (herein referred to as Rac1^Dead^; **Figure S1A**), could not promote Arf1 binding at the same concentration (**Figure 1C**, blue). Thus, Rac1 can enhance the weak interaction between Arf1 and WRC by ∼30 fold.

We found Arf1 binding was likely mediated by the Sra1 or Nap1 subunit, but not WAVE1, Abi2 or HSPC300, as only the dimeric subcomplex containing Sra1/Nap1, but not the trimeric subcomplex formed by WAVE1/Abi2/HSPC300, showed weak binding signals comparable to the fully assembled, pentameric WRC (**Figure 1D**, lane 4 vs. 6, asterisks). Unlike binding to the intact WRC, however, the interaction with the Sra1/Nap1 dimer could not be enhanced by Rac1 (**Figure 1D**, lane 5 vs. 7, asterisks), suggesting that even though Sra1 or Nap1 may contain the Arf1 binding site, the enhancement of Arf1 binding by Rac1 is dependent on the fully assembled WRC. Moreover, we found that Arf1 binding to WRC was sensitive to both pH and salt concentration, with pH 6-7 and 50 mM NaCl, but not pH 8 or 100 mM NaCl being able to sustain the binding (**Figure 1E**, lane 2, 6). This indicates that the Arf-WRC binding involves polar interactions (see below).

We further tested if the Arf1-WRC interaction is unique to Arf1 or is general to other Arf-family proteins. In mammals, the Arf family contains six canonical members (Arf1-Arf6) and various distantly related Arf-like proteins (Arl) (Gillingham and Munro, 2007; Sztul et al., 2019). Based on sequence similarities, Arf1-Arf6 can be further divided into three classes: Class I (Arf1, 2, 3), Class II (Arf4, 5) and Class III (Arf6). We found that besides Arf1, Arf5 and Arf6 also robustly bound the WRC in a Rac1-dependent manner, although perhaps with slightly different affinities (**Figure 1F**, lane 1-6). By contrast, Arl1 or Arl2 did not show clear binding (**Figure 1F**, lane 7-10). These results suggest that the six members of Arf family, but perhaps not the more divergent Arl proteins, can use the same mechanism to interact with the WRC. Together, our biochemical reconstitution established a direct, nucleotide-dependent interaction between Arf-family GTPases and WRC. This interaction is greatly enhanced by Rac1 binding to the WRC.

### Arf1 binding mainly depends on Rac1 binding to the D site

Rac1 can bind to both the A and D sites on WRC, albeit with distinct affinities and effects on WRC activation (Chen et al., 2017; Ding et al., 2022). Therefore, we asked which Rac1 binding event was key to promoting Arf1 binding. To answer this question, we first used single amino acid mutations to specifically disrupt the A or D site from binding to Rac1 (Chen et al., 2017, 2010; Ding et al., 2022) (**Figure 2A**, cartoon). When Rac1 binding to the A site was disrupted by Sra1^C179R^, Rac1 binding to the D site still enhanced Arf1 binding, but to a lower extent than the WT WRC (**Figure 2A**, lane 6, 7). By contrast, when Rac1 binding to the D site was disrupted by Sra1^Y967A^, Rac1 binding to the A site could no longer promote Arf1 binding (**Figure 2A**, lane 8, 9). These data indicate that Rac1 binding to the D site plays a more important role in promoting Arf1 binding.

**Figure 2.**
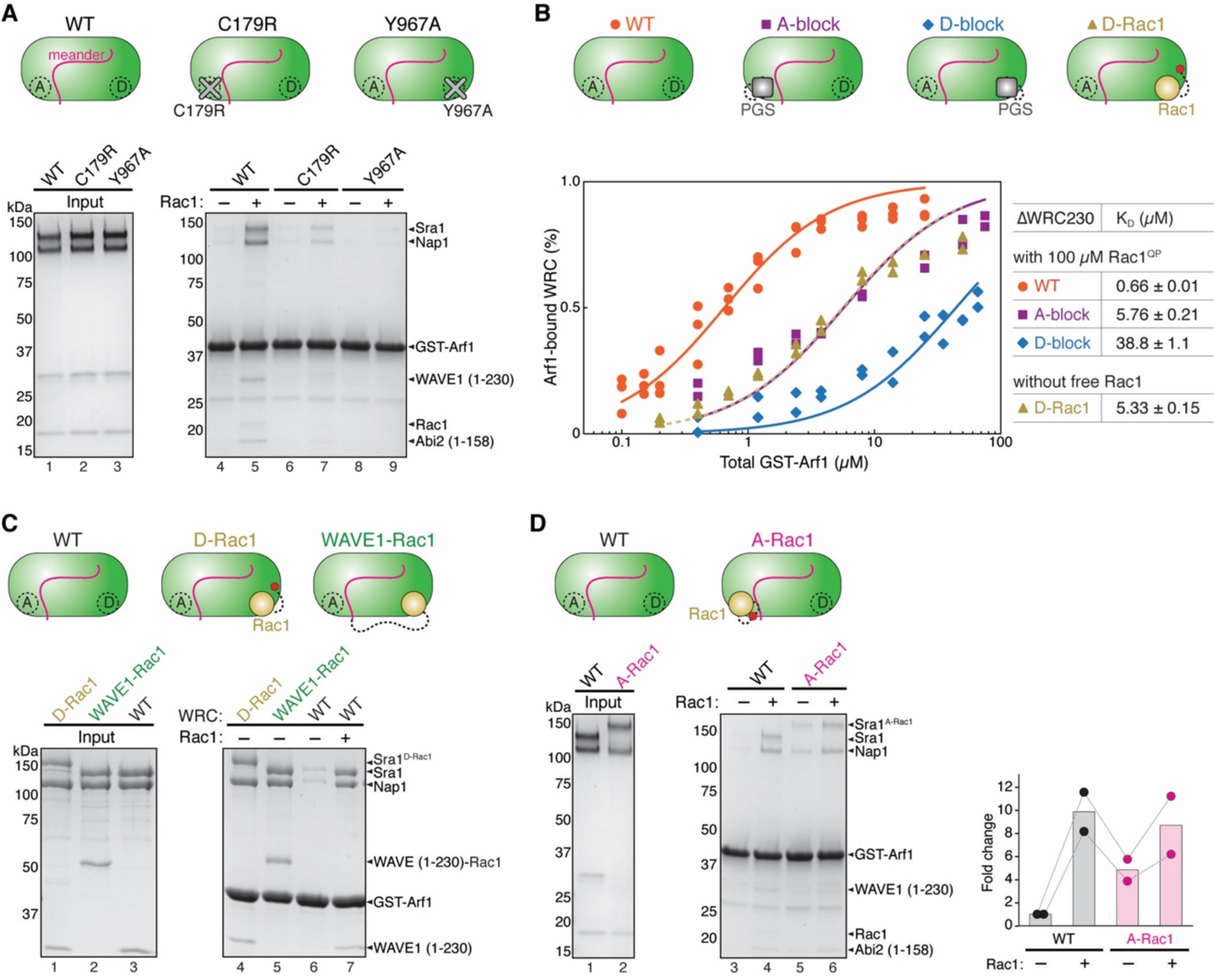
Arf1 binding mainly depends on Rac1 binding to the D site. **(A)** Coomassie blue-stained SDS-PAGE gels showing GST-Arf1 pull-down of WRC bearing point mutations in Sra1 that specifically disrupt the A or D site. **(B)** EPD assay measuring the binding affinity of GST-Arf1 for the indicated ΔWRC230 constructs in the presence or absence of 100 µM Rac1^QP^. Data for each mutant are pooled from two independent experiments. Data for the WT WRC are taken from Figure 1C and used here as a reference point. See Figure S1 for representative gel images. **(C)** Coomassie blue-stained SDS-PAGE gels showing GST-Arf1 pull-downs of WRCs with Rac1 tethered to indicated positions to stabilize Rac1 binding to the D site. **(D)** Coomassie blue-stained SDS-PAGE gels showing GST-Arf1 pull-down of WRCs with Rac1 inserted between Y423/S424 of the surface loop (a.a. 418-432) to stabilize Rac1 binding to the A site. Shown on the right is the gel quantification of the Sra1-Nap1 bands normalized to GST-Arf1 bands from two independent repeats, with the data from each repeat connected. In the schematic of WRCs, red dots indicate the tethering points of Rac1 to A or D site.

To further validate this result, we used EPD assays to directly measure the binding affinities of Arf1 to WRCs with disrupted A vs. D site. For this, instead of using the above single amino acid mutations to disrupt either site, which may retain weak, residual Rac1-binding activity, we inserted an inert protein PGS (glycogen synthase from the extreme thermophile *Pyrococcus abysii*) to a surface loop at the A or D site to completely block Rac1 binding. We herein name the new variants WRC^A-block^ and WRC^D-block^, respectively (**Figure 2B**, cartoon). Being a small, stable protein and with its N- and C-termini located in close proximity (6.5 Å), PGS was initially used to insert into the human orexin/hypocretin receptors hOX1R and hOX2R to stabilize an intracellular loop and produce high-resolution diffracting crystals (Yin et al., 2015, 2016). Inserting PGS into the surface loop of the A or D site did not affect WRC assembly or purification (**Figure S2A, B**) or the basal level of Arf1-WRC interaction in the absence of free Rac1 (**Figure S1C**), but indeed further reduced the affinity measurement of Rac1 to WRC (from K_D_ ∼ 2 µM for WRC^Y967A^ to ∼7.5 µM for WRC^D-block^; **Figure S1B**, blue vs. orange), likely due to eliminating the residual Rac1 binding to the D site in WRC^Y967A^. When we blocked the A site and subjected the D site to 100 µM Rac1, Arf1 binding was enhanced, although not to the level of WT WRC (K_D_ ∼5.76 µM for WRC^A-block^ vs. ∼0.66 µM for the WT WRC; **Figure 2B**, purple vs. orange; **Figure S1D**), suggesting Rac1 binding to the D site was partially sufficient to promote Arf1 binding. By contrast, when we blocked the D site and exposed the A site to 100 µM Rac1, Arf1 binding was not enhanced, but remained similar to that in the absence of Rac1 or in the presence of 100 µM Rac1^Dead^ (K_D_ ∼38.8 µM; **Figure 2B**, blue; **Figure 1C**, black; **Figure S1C**, orange), suggesting Rac1 binding to the A site alone could not promote Arf1 binding in this specific experimental condition (but see below).

As an alternative strategy to validate the contribution of the D site to Arf1 binding, we stabilized Rac1 binding to the D site by tethering it to the C-terminus of Sra1 (which we refer to as ΔWRC230^D-Rac1^) (Ding et al., 2022) or the C-terminus of WAVE1 that lacked the WCA (which we named ΔWRC230^WAVE1-Rac1^) (Chen et al., 2017) (**Figure 2C**, cartoon). These constructs stabilize D site Rac1 binding, which had allowed us to solve cryo-EM structures of the WRC with Rac1 bound to the D site (Chen et al., 2017; Ding et al., 2022). We found that, without free Rac1, both ΔWRC230^D-Rac1^ and ΔWRC230^WAVE1-Rac1^ were able to enhance Arf1 binding to the level of the WT WRC enhanced by free Rac1 (**Figure 2C**, lane 4, 5, 7). Furthermore, in the EPD assay, ΔWRC230^D-Rac1^ without free Rac1 enhanced Arf1 binding to a level nearly identical to that of the WRC^A-block^ in the presence of 100 µM Rac1 (K_D_ ∼5.33µM; **Figure 2B**, golden vs. purple). Therefore, supplying Rac1 to the D site by covalent tethering has the same effect in promoting Arf1 binding as supplying free Rac1 to a WRC with a blocked A site.

The above assays confirm that Rac1 binding to the D site is essential for enhancing Arf1 binding, but also show mutating or blocking the A site dampens this effect (**Figure 2A**, lane 7; **Figure 2B**, purple). This indicates that Rac1 binding to the A site should also play a role, which might have eluded detection in the assays described above due to the low affinity of the A site for Rac1. The potential cooperativity between A and D sites could further reduce A site binding when the D site is disrupted (Chen et al., 2017; Ding et al., 2022). To examine the contribution of the A site more directly, we stabilized Rac1 binding to the A site by inserting a Rac1 between Y423/S424 in a non-conserved surface loop near the A site (termed ΔWRC230^A-Rac1^; **Figure 2D**, cartoon). This strategy had allowed us to determine the cryo-EM structure of the WRC with Rac1 bound to the A site and D site simultaneously (Ding et al., 2022). We found that, without free Rac1, tethering Rac1 to the A site mildly promoted Arf1 binding (**Figure 2D**, lane 3 vs. 5). Adding free Rac1 to ΔWRC230^A-Rac1^ to occupy the D site further enhanced Arf1 binding (**Figure 2D**, lane 6). These data suggest Rac1 binding to the A site partially contributes to Arf1 binding. Taken together, we conclude Rac1 binding to both A and D sites plays a role in promoting Arf1 binding to the WRC, but with the D site having a major contribution as compared to the A site.

### Arf1 promotes WRC activation using a novel mechanism distinct from Rac1

Arf1 and Rac1 were shown to cooperatively promote WRC activation on lipid-coated beads (Koronakis et al., 2011). Since Rac1 binding to the A site is sufficient to activate the WRC through an allosteric mechanism (Ding et al., 2022), the question remains: does Arf1 binding merely increase the membrane recruitment of WRC, contribute to the same allosteric changes driven by Rac1 binding to the A site, or promote WRC activation through an entirely different mechanism?

To distinguish between these possibilities, we first tested if Arf1 differentially binds to the WRC in the autoinhibited (“closed”) or activated (“open”) state. Previous studies showed that, as an activator, Rac1 had higher affinity for the “open” conformation represented by ΔWRC230 (which lacks the WCA) than for the “closed” conformation represented by the WRC that contained WCA (WRC230WCA; **Figure 3A**, cartoon) (Chen et al., 2017, 2010). If Arf1 is an activator, it should similarly prefer the “open” conformation. Indeed, we observed less binding for WRC230WCA than ΔWRC230, both in the presence and absence of Rac1 (**Figure 3A**). Our EPD assay further confirmed this observation (**Figures 3B** and **S1E**). In the absence of free Rac1, Arf1 had very low binding affinity for WRC230WCA, with a K_D_ (∼107 µM) ∼5 times of ΔWRC230 (∼22.6 µM) (**Figure 3B**, blue vs. black). Addition of a saturating concentration of Rac1 (100 µM) enhanced Arf1 binding to both WRC230WCA and ΔWRC230, although not to the same level (K_D_ ∼8.2 µM for WRC230WCA vs. K_D_ ∼0.66 µM for ΔWRC230) (**Figure 3B**, purple vs. orange). These data indicate Arf1 distinguishes the “closed” vs. the “open” conformation and therefore may act as an activator of the WRC.

**Figure 3.**
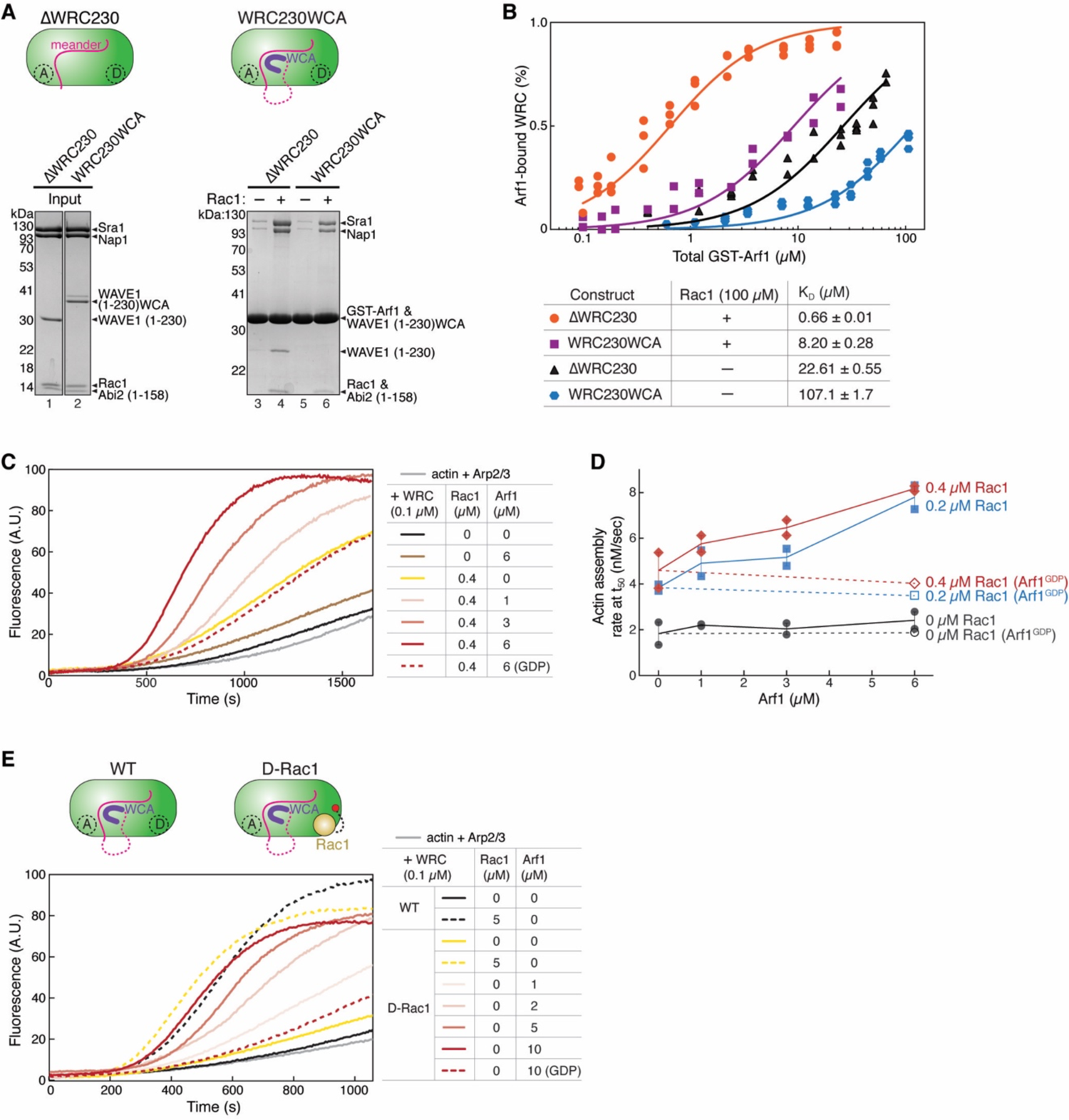
Arf1 promotes WRC activation independent of Rac1 binding to the A site. **(A)** Coomassie blue-stained SDS-PAGE gels showing GST-Arf1 pull-down of WRC with or without the WCA sequence. **(B)** EPD assay comparing the binding affinity of GST-Arf1 for the WRC with or without the WCA sequence. Data for WRC230WCA are pooled from two independent experiments for each condition. Data for the ΔWRC230 are taken from Figure 1C and used here as a reference point. See Figure S1 for representative gel images. **(C-D)** Representative pyrene-actin polymerization assay (C) and quantification of the actin polymerization rate at t_50_ (D) (Doolittle et al., 2013) measuring activity of WRC230WCA in the presence of indicated concentrations of Rac1^QP^ and Arf1. **(E)** Pyrene-actin polymerization assay of the WT WRC230WCA vs. WRC^D-Rac1^ in response to the addition of free Rac1^QP^ or Arf1. Reactions in (C-E) contain 3.5 µM actin (5% pyrene labeled), 10 nM Arp2/3 complex, 100 nM WRC, and indicated amounts of Rac1 and/or Arf1. In all assays, Arf1 is loaded with GMPPNP, unless it is indicated with GDP.

We next measured whether Arf1 could promote WRC activation in the pyrene-actin polymerization assay in aqueous solution (as opposed to on membranes as in the previous study (Koronakis et al., 2011)) (**Figure 3C-E**). For this, the Arf1 construct used in this study does not contain the N-terminal amphipathic helix (also referred to as Arf1^ΔN17^). This helix is important for Arf1 to bind membranes, but is usually dispensable for binding downstream effectors and therefore often removed in biochemical and structural studies (Dong et al., 2012; Ren et al., 2013). In the absence of Rac1, Arf1 had no obvious effect on WRC activity, potentially due to its low binding affinity to WRC230WCA (**Figure 3C**, brown; **3D**, black). In the presence of low concentrations of Rac1, however, Arf1 enhanced WRC activation in a dose-dependent manner (**Figure 3C**, red curves; **3D**, red, blue). The enhanced WRC activation depended on Arf1 GTP-binding as Arf1 loaded with GDP had no such an effect (**Figure 3C-D**, dashed lines).

The above data suggest that, in addition to potentially recruiting WRC to the plasma membrane, Arf1 directly contributes to WRC activation. Due to the presence of free Rac1 in the above reactions, however, these data cannot tell whether Arf1 acts by promoting the same conformational changes driven by Rac1 binding to the A site, or by directly activating the WRC through a separate mechanism. To distinguish between these two mechanisms, we further tested if Arf1 could activate the WRC230WCA in which a Rac1 molecule was tethered to the D site (WRC^D-Rac1^; **Figure 3E**, cartoon) (Ding et al., 2022). In this construct, the tethered Rac1 does not activate the WRC (Ding et al., 2022) (also see **Figure 3E**, yellow solid curve), but can promote Arf1 binding to the WRC (**Figure 2C**), allowing us to determine whether Arf1 can activate WRC in the absence of a Rac1 molecule acting through the A site. Remarkably, in the absence of free Rac1, we found Arf1 activated WRC^D-Rac1^ in a dose-dependent manner (**Figure 3E**, red solid curves), while Arf1 loaded with GDP had no such effect (**Figure 3E**, red dotted curve). To rule out the possibility that Arf1 may activate WRC by mimicking Rac1 binding to the A site, we disrupted the A site by the point mutation C179R and found Arf1 still activated WRC^D-Rac1^ in a dose-dependent manner (**Figure S3A**), although with reduced potency perhaps because the mutation indirectly weakened Arf1 binding. Together, the above data suggest that Arf1 binding can directly activate WRC, at least *in vitro*, and the Arf1-mediated activation does not involve an interaction of either Rac1 or Arf1 with the A site. Therefore, Arf1 must use a novel mechanism to drive WRC activation.

It is important to note that the Arf1-mediated WRC activation reached levels similar to those achieved with Rac1 binding to the A site (**Figure 3E**, black and yellow dotted curves), suggesting Arf1 binding activates the WRC by releasing the WCA, instead of by causing protein aggregation (which is believed to cause artificial WRC activation to a much larger extent than the release of WCA) (Eden et al., 2002; Gautreau et al., 2004; Lebensohn and Kirschner, 2009). This is consistent with our dynamic light scattering (DLS) measurement showing that Arf1 did not promote WRC aggregation (**Figure S2J**).

### Arf1 binds to a conserved site distinct from Rac1 binding sites

How does Arf1 binding activate WRC? To answer this question, we determined the Arf1 binding site by combining protein docking, surface conservation analysis, mutagenesis, and molecular dynamics simulation (**Figures 4, 5, S4, S5**). We first searched for potential binding sites by using several different protein docking programs, including ClusPro (Desta et al., 2020), HADDOCK (Van Zundert et al., 2016), InterEvDock (Quignot et al., 2018), FRODOCK (Ramírez-Aportela et al., 2016), and HDOCK (Yan et al., 2020). During the search, we restrained the Switch I and Switch II motifs of Arf1 in close contact with the WRC, since they usually mediate GTPase-effector interactions. Combining the docking results with the surface conservation analysis of the WRC by Consurf (Ashkenazy et al., 2016), we selected a series of conserved surface patches, mutated the solvent-exposed residues individually or in combination, purified the mutant WRCs, and used pull-down assays to examine if any mutations could disrupt Arf1 binding (**Figure S4**).

**Figure 4.**
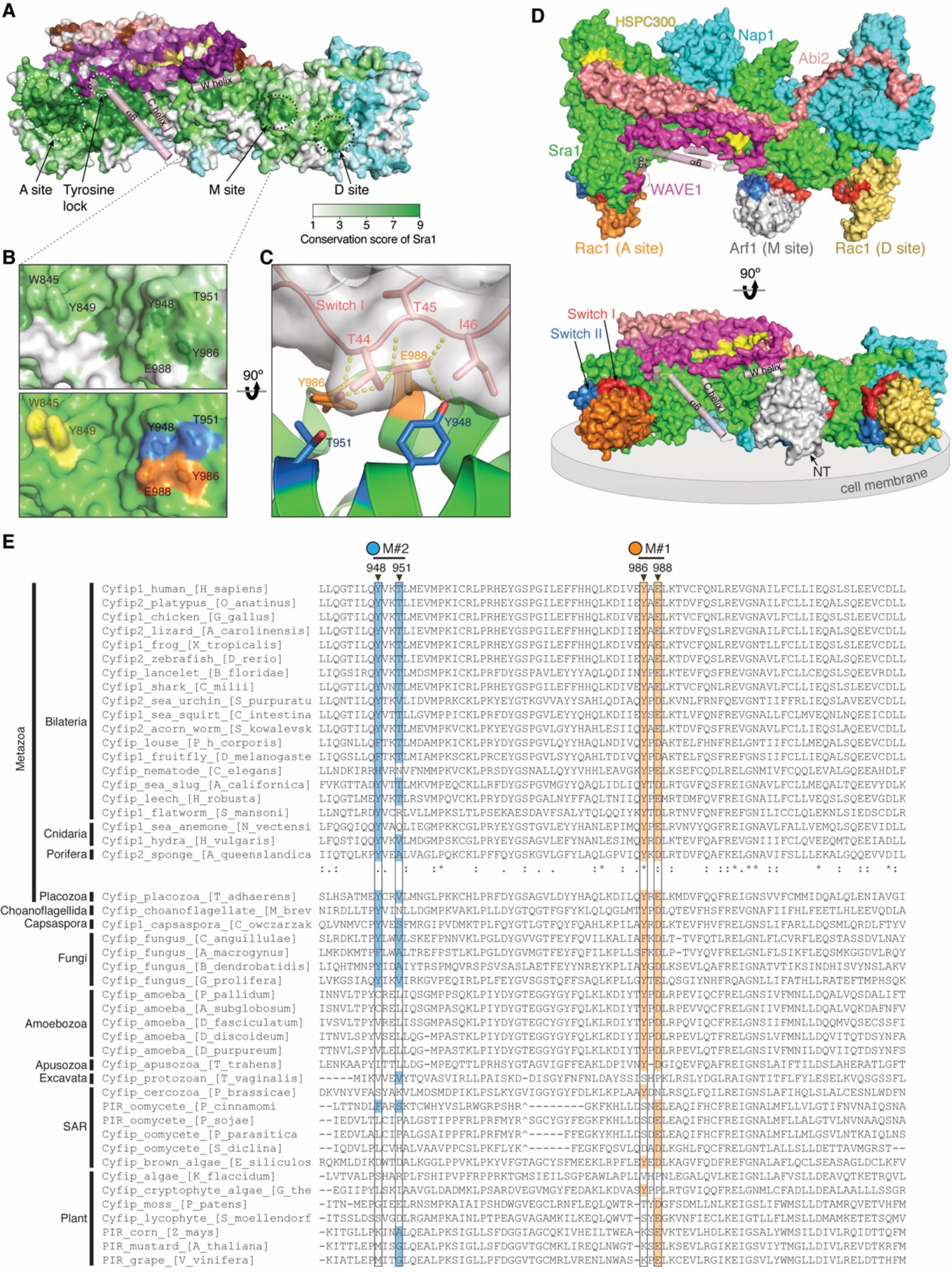
Arf1 binds to a conserved site distinct from Rac1 binding sites. **(A)** Surface conservation of the WRC, with color to white gradients representing the most conserved surface residues (ConSurf score = 9 for darkest colors) to the least conserved residues (ConSurf score = 1 for white color) (Ashkenazy et al., 2016). Important sites on Sra1, including the identified Arf1 binding site (M site), are indicated with dotted circles. Semitransparent pink cylinders refer to the sequence in WAVE1 that are destabilized upon WRC activation by Rac1 (Ding et al., 2022). **(B)** Close-up view of the M site showing surface conservation (top) and surface patches to be mutated (bottom, colored using the same scheme in Figure 5A). **(C)** Side view showing the interaction between Arf1 and the M site in the MD-optimized model C8. Contacting residues are shown as sticks. Yellow dotted lines indicate polar interactions. **(D)** Surface representation of the overall structural model of the WRC bound to two Rac1 molecules (PDB: 7USE) (Ding et al., 2022) and one Arf1 molecule (PDB: 1J2J). Position of Arf1 shows the MD-optimized docking solution C8, which has the lowest binding free energy. Switch I and II elements of Rac1 and Arf1 GTPases are red and blue, respectively. Grey disc demonstrates the predicted orientation of the WRC at the inner surface of the plasma membrane. The N-terminus of Arf1^ΔN17^ used in this study is indicated with arrow. **(E)** Sequence alignments of Sra1 from representative eukaryotic organisms. Surface residues of the M site (black boxes) are highlighted with orange for the M#1 surface patch and blue for M#2, as indicated by black arrowheads on top. Degrees of conservation in animals (up to Porifera) are represented with ClustalW symbols (Thompson et al., 1994)(* for no change,: for conserved,. for less conserved changes). ‘-’ for missing amino acids; ‘^’ for amino acid insertions in alignments that were not shown.

**Figure 5.**
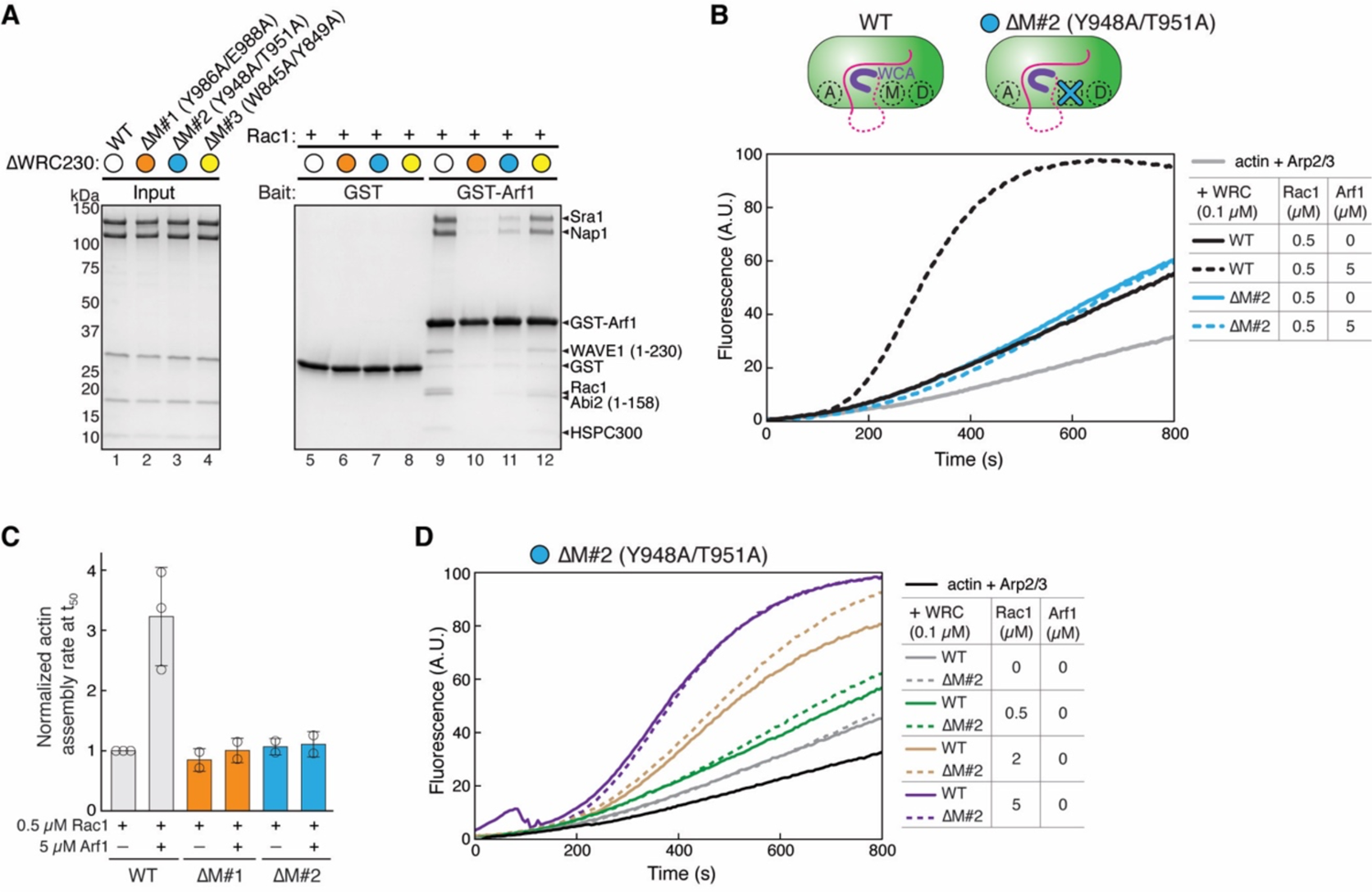
M site mutations disrupt Arf1 binding and Arf1-mediated WRC activation. **(A)** Coomassie blue-stained SDS-PAGE gels showing GST pull-down of ΔWRC230 bearing the indicated mutations in Sra1 at the M site. **(B-C)** Representative pyrene-actin polymerization assay (B) and quantification of the actin polymerization rate at t_50_ normalized to WT WRC230WCA + 0.5 µM Rac1 (C), measuring the effect of M site mutations on WRC activation by Arf1. Reactions contain 3.5 µM actin (5% pyrene labeled), 10 nM Arp2/3 complex, 100 nM WRC230WCA (WT or indicated mutants), and indicated amounts of Rac1^QP^ and/or Arf1 loaded with GMPPNP. Error bars represent standard errors of means. **(D)** Comparison of the WT WRC to the ΔM#2 (Y948A/T951A) mutant activated by different amounts of Rac1^QP^. Reactions were performed in the same conditions as in (B).

Out of over 12 conserved surfaces that we surveyed, one surface specifically disrupted Arf1 binding (**Figure S4A**, M1 site; **S4F,** lane 6; **Figure 5A**, lane 10). We named this site the M site since it is in the **<u>m</u>**iddle of the WRC, sandwiched between the D site and the W helix of the WCA (**Figure 4A**). The M site is a small, conserved, and slightly negatively charged surface patch on Sra1 (**Figure S4A, B**). Mutating the conserved surface residues at the M site, either Y986A/E988A (ΔM#1) or Y948A/T951A (ΔM#2), disrupted Arf1 binding, whereas mutating two other conserved residues, W845A/Y849A (ΔM#3), near the M site did not disrupt Arf1 binding (**Figure 4B**; **Figure 5A**, lane 10-12; **Figure S4F**), suggesting the effect of ΔM#1 and ΔM#2 was specific to Arf1 binding. Furthermore, the WRC carrying ΔM#1 or ΔM#2 mutations could not be further activated by Arf1 (**Figures 5B, C** and **S3B**). It’s important to note that the M site mutations only disrupted Arf1-mediated activation, but not Rac1-mediated activation (**Figures 5D** and **S3B**). Thus, these surface mutations are specific in disrupting Arf1 binding and Arf1-mediated activation, without affecting WRC folding (**Figure S2C-G**) or disturbing Rac1-mediated activation.

Note all four residues are highly conserved in animals (**Figure 4E**). In particular, Y986 remains strictly Tyrosine from human to sponge, while E988 is only exchangeable with Aspartate. In non-animal species they are either partially conserved (such as in amoeba) or not conserved (such as in plants) (**Figure 4E**). This suggests the Arf-WRC interaction is important for processes unique to animals.

To further define the binding mechanism, we applied molecular dynamics (MD) simulation to optimize binding poses of the top 6 docking models that placed Arf1 at the M site (**Figure S5A, B**). We then evaluated different models by calculating the molecular mechanics/Poisson–Boltzmann surface area/weighted solvent accessible surface area (MM-PBSA-WSAS) free energies of the whole complex and the binding free energy between Arf1 and WRC (**Figure S5C-I**). Out of the 6 docking models, model C8 gave the lowest binding free energy (**Figure S5G, I**). Importantly, introducing ΔM#1 or ΔM#2 mutations onto model C8 increased the binding energy, suggesting they destabilized Arf1-WRC interaction. By contrast, introducing the control mutation ΔM#3 did not affect the binding energy (**Figure S5I**). These data are consistent with our pull-down assays showing only ΔM#1 & 2, but not ΔM#3, disrupted Arf1 binding (**Figure 5A, S4F**).

Note that it was previously shown the M371V mutation in Hem1 (M373V in Nap1) found in human patients interfered with (but did not abolish) Arf1 binding and WRC activation (Cook et al., 2020). The above analysis suggests M371 is not the Arf1 binding site. Rather, the effect of M371V was likely indirect, as this residue is located at the bottom of a deep pocket neighboring the D site, where it was difficult to accommodate an Arf1 molecule (**Figure S4A**).

The MD-optimized model sheds light on how Arf1 binds and activates WRC. First, the interaction is mediated by the Switch I motif (**Figures 4C** and **S5G**), the same region that binds to EspG (Dong et al., 2012), explaining how EspG competes off WRC binding to inhibit phagocytosis during pathogenic *E. coli* infection (Humphreys et al., 2016) (**Figure 1B**). Second, the interaction mainly involves hydrogen bonding between Y986 and E988 in Sra1 and T44 and T45 in Arf1, with Y948 or T951 in Sra1 contacting I46 and T44 in Rac1 through Van der Waals interactions (**Figures 4C** and **S5G**). This explains why Y986A/E988A (ΔM#1) disrupted Arf1 binding more severely than Y948A/T951A (ΔM#2) in GST pull-down assays (**Figures 5A** and **S4F**), and is also consistent with our observation that Arf1 binding is sensitive to pH and salt concentration (**Figure 1E**). Third, the relative orientation of Arf1 is compatible with the model of how WRC is oriented on the membrane together with two Rac1 molecules (**Figure 4D**) (Chen et al., 2017, 2010; Ding et al., 2022). In this orientation, the N-terminus of Arf1^Δ17^ is near the plasma membrane (**Figure 4D**, arrow), which would allow its N-terminal amphipathic helix to associate with membranes. Finally, this model explains how Arf1 binding may activate the WRC. Arf1 is located near (but not in direct contact with) the W helix of WCA (**Figure 4D**). Therefore, distinct from Rac1-mediated WRC activation, which involves a series of conformational changes propagating from the A site to a conserved region around WAVE1^Y151^ (referred to as Tyrosine lock) to release the WCA (Ding et al., 2022), Arf1 binding may contribute to WRC activation by directly perturbing the W helix located in its proximity (see models in **Figure 7**).

The identification of the M site allowed us to specifically probe the function of the Arf1-WRC interaction in the cell. WRC is key to actin polymerization at plasma membranes and formation of sheet-like protrusions known as lamellipodia and membrane ruffles commonly found at the leading edge of migrating or spreading cells (Rottner et al., 2021; Takenawa and Suetsugu, 2007) (**Figure 6A**). In our previous complementation assays using B16-F1 cells genetically disrupted for both *Sra1* and *Cyfip2* genes, mutating the A site almost completely abolished WRC-mediated lamellipodia formation, while mutating the D site impaired (but did not eliminate) actin assembly and lamellipodia morphology (Schaks et al., 2018, 2020). Using the same assay, we found mutating the M site produced phenotypes nearly identical to mutating the D site (**Figure 6B-D**). In both cases, mutations led to narrow actin networks and reduced the formation frequency of mature lamellipodia, but without affecting WRC localization or assembly (**Figure 6B, D)**. Interestingly, when we combined the M site and D site mutations into one construct, they did not aggravate the phenotype, except that the ΔD/ΔM#2 dual mutations slightly decreased the total percentage of lamellipodia-containing cells (**Figure 6E-G**). Together, the above results suggest that the M and D sites act in the same mechanistic pathway to regulate lamellipodial morphology, with Arf1 binding to the M site likely acting downstream of Rac1 binding to the D site (see *in vitro* data above).

**Figure 6.**
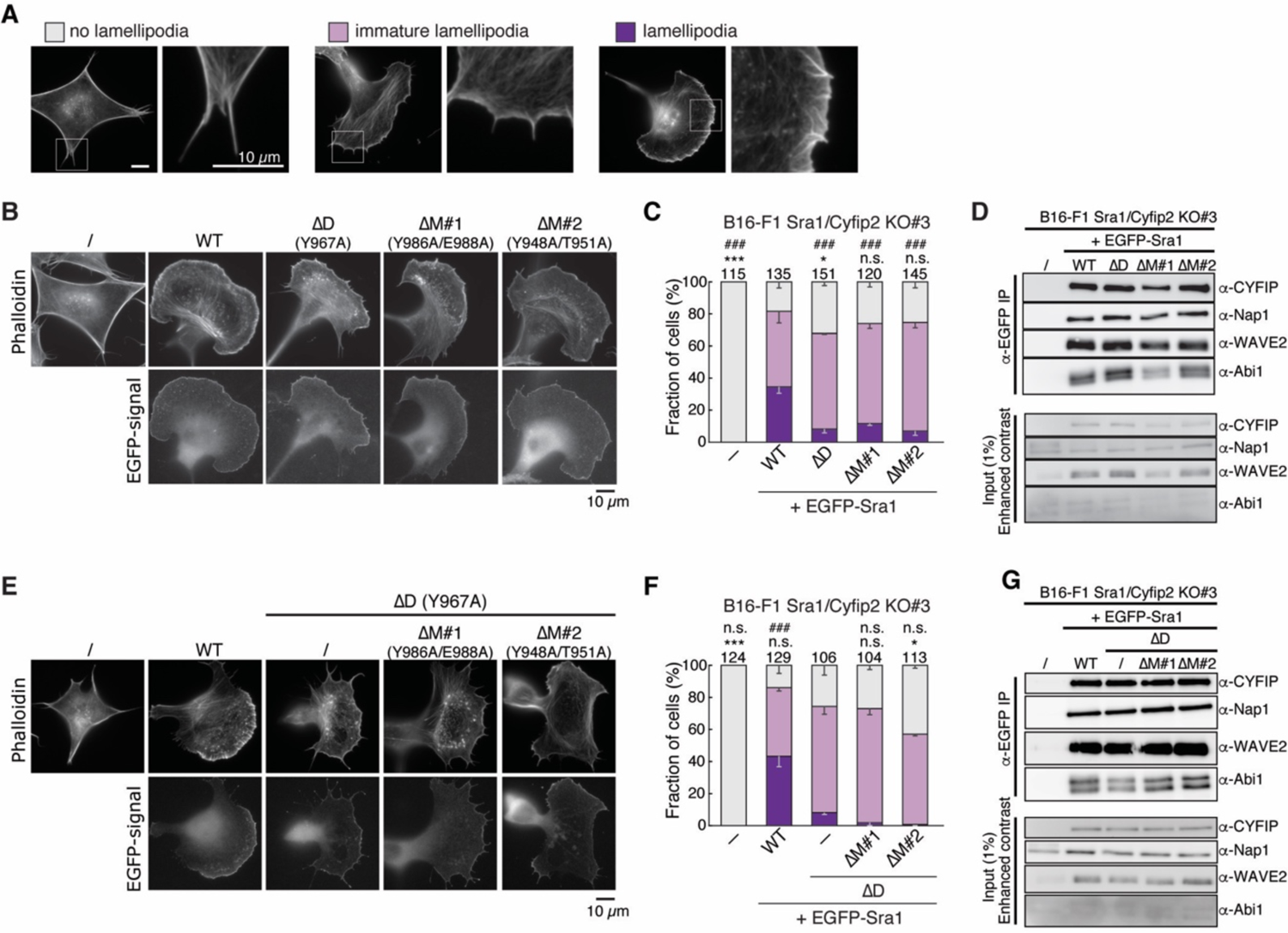
M site mutations impaired lamellipodia morphology. **(A, B, E)** Representative fluorescence images of lamellipodia formation in B16-F1 *Sra1/Cyfip2* double KO#3 cells transfected with indicated EGFP-Sra1 variants and stained by phalloidin for F-actin. Images show representative examples of the cell morphologies used for categorization of induced effects below. ΔD for Sra1^Y967A^, which disrupts the D site. ΔM #1 for Sra1^Y986A/E988A^. ΔM #2 for Sra1^Y948A/T951A^. **(C, F)** Quantification of lamellipodial morphologies. Statistical significance was assessed from 3 repeats for differences between cells transfected with WT (wild type) (**C**) or ΔD (**F**) vs. no (-) or indicated constructs concerning cell percentages displaying “no lamellipodia” phenotype (* p < 0.05; *** p < 0.001) and with “lamellipodia” phenotype (### p < 0.001). n.s.: not statistically significant. Error bars represent standard errors of means. Numbers of cells used in the quantification are shown on top of each column. **(D, G)** Immunoprecipitation (IP) and Western blot of the same B16-F1 *Sra1/CYFIP2* double KO#3 cells used in (B & C and E & F), respectively. The cells were transfected with indicated EGFP-tagged Sra1 variants, lysed, and probed for the expression and assembly of WRC, as exemplified by CYFIP (for both Sra1 and Cyfip2), Nap1, WAVE2, and Abi1.

**Figure 7.**
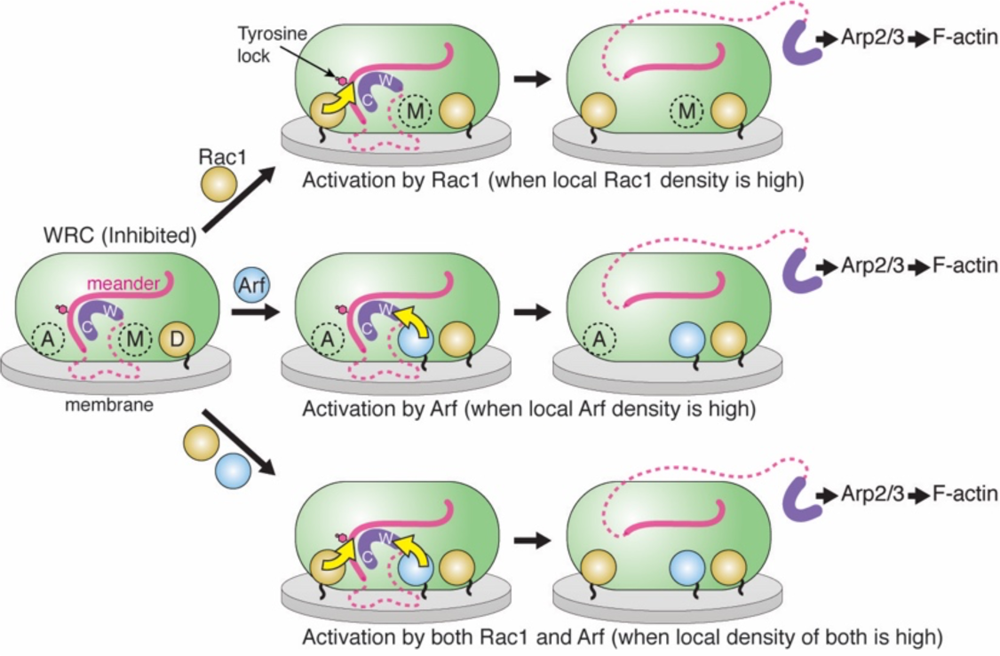
Rac1 and Arf may act both cooperatively and separately to promote WRC activation. Schematic showing how the WRC can be activated by Rac1 (top), Arf (middle) and both (bottom) through specific mechanisms that can arise independently from each other. Structural elements critical to WRC inhibition and activation are shown. Yellow arrows indicate structural pathways leading to WRC activation. Magenta dashed lines represent unstructured sequences in WAVE1. Black wiggly lines attached to Arf and Rac1 represent membrane binding sequences and lipid modifications of the GTPases. Rac1 first engages with the D site due to its relatively high affinity, which primes the WRC on the membrane without causing activation (left). When Rac1 density on the membrane is high (top), further binding of Rac1 to the A site promotes WRC activation by allosterically destabilizing the Tyrosine lock region, which subsequently releases Y151 (indicated by pink hexagon) and the WCA (purple) (Ding et al., 2022). Alternatively, when Arf1 density on the membrane is high (middle), Rac1 at the D site promotes Arf binding to the M site, which in turn, through its close proximity to the W helix, can perturb WCA binding to promote WRC activation. The remaining part of the schematic displays the functional outcome of both mechanisms operating in cooperation to ensure an optimized output response (bottom).

## Discussion

By biochemical reconstitution and structural analysis, our work establishes that Arf1 directly interacts with the WRC through a previously unidentified conserved surface located on Sra1. We show that, although intrinsically weak, this interaction can be greatly enhanced by Rac1 binding to the D site. Once bound to the WRC, Arf1 can independently drive WRC activation, at least *in vitro*, by using a mechanism distinct from that mediated by Rac1 binding to the A site. We further demonstrate that disrupting Arf1-WRC interaction by point mutations specifically abolishes Arf1-mediated (but not Rac1-mediated) WRC activation, and impairs WRC-mediated lamellipodia formation. Our work has important implications on the regulation of the actin cytoskeleton in many different biological systems.

First, our study established a new mechanism underlying WRC activation. The WRC is a central signaling hub through which a large diversity of membrane ligands can transmit signals to Arp2/3 complex-mediated actin polymerization (Chen et al., 2014b, 2017; Lebensohn and Kirschner, 2009; Rottner et al., 2021). Despite the long list of WRC ligands, Rac1 has been known as the only activator that is both necessary and sufficient— at least *in vitro*—to activate WRC (Chen et al., 2017; Schaks et al., 2018). While other ligands may act cooperatively with Rac1 to further tune WRC activity, exactly how they do so is completely unknown (Chen et al., 2014b, 2014c; Koronakis et al., 2011; Lebensohn and Kirschner, 2009). In particular, how Arf1 facilitates WRC activation has remained enigmatic for many years. It was not known if Arf1 can directly interact with WRC, and if yes, how Arf works together with Rac1 to promote WRC activity (Singh et al., 2019). Our work provides firm answers to these questions, revealing that significant Arf1 binding relies on Rac1 binding mainly to the D site, but Arf1 binding can directly promote WRC activity even independently of Rac1 binding the A site. These results establish Arf1 as a second, genuine activator of the WRC and provides a mechanism to explain the cooperativity between Arf1 and Rac1 previously observed both *in vitro* and in cells (Boshans et al., 2000; Koronakis et al., 2011; Radhakrishna et al., 1999; Santy and Casanova, 2001).

Second, our study lays a foundation for studying how WRC-mediated actin polymerization connects various Arf- and Rac1-mediated processes. Our work identifies point mutations that can specifically disrupt Arf binding and Arf-mediated (but not Rac1-mediated) WRC activation. These mutations will be powerful tools for dissecting the role of the Arf-WRC-Arp2/3-actin signaling axis from the canonical Rac1-WRC-Arp2/3-actin axis. Arf-family GTPases play an important role in various membrane trafficking processes, with some of them tightly connected to actin cytoskeleton regulation (Myers and Casanova, 2008; Singh et al., 2017; Sztul et al., 2019). On the other hand, new roles of actin, WRC, and Arp2/3 complex are emerging, suggesting their importance in the endomembrane systems beyond their canonical role in driving plasma membrane protrusions (Anitei et al., 2010; Cheng et al., 2007; Kang et al., 2010; Sung et al., 2008). We thus posit that Arf-mediated WRC activation provides the cell with an additional pathway for promoting WRC activation and actin polymerization, the precise outcome of which will likely depend on relative local membrane densities of Rac1 vs. Arf (**Figure 7**). Specifically, Rac1 binding to the high-affinity D site may serve as a general recruitment mechanism to prime the WRC on the membrane without causing activation. Then, depending on specific upstream signals in distinct cell types and tissues leading to activation of various Arf- or Rac1-GEFs, the precise tuning of WRC activation in given condition and system will depend on the local density of activated Rac1 or Arf molecules, which can subsequently trigger WRC activation by distinct structural mechanisms (**Figure 7**).

Third, the Arf binding site is highly conserved in animals, from human to sponge, but is only partially conserved in other organisms and is not conserved in plants. This suggests that the function of Arf1-mediated WRC activation is likely important for processes unique to animals, such as neuronal outgrowth and synapse formation, immune cell chemotaxis and activation, and cancer cell migration and metastasis, in all of which Arf and WRC play important roles (Donaldson and Jackson, 2011; Myers and Casanova, 2008; Rottner et al., 2021; Singh et al., 2017; Sztul et al., 2019). In non-animal species, while sequence analysis of the M site suggests the direct interaction between Arf and the WRC is perhaps lost (**Figure 4E**), considering the conserved importance of Arf and WRC in non-animal species, we cannot rule out the possibility that the M site surface and Arf molecules may still have co-evolved to keep the connection between Arf and the WRC maintained. Our work raises the possibility of exploring the role of Arf-related processes in WRC-mediated actin polymerization in both animal and non-animal organisms.

Together, this work uncovers a new, conserved mechanism underlying WRC activation, and provides a foundation for exploring the regulation of the actin cytoskeleton in multiple processes in which Rac and the various Arf-family GTPases may intimately cooperate.

## Acknowledgements

We thank Shae Padrick at Drexel University for updating the published python codes for actin data analysis, Aubrey Sijo-Gonzales, Finlan Rhodes, Leyuan Loh, Ganesh Prasad, Simanta Mitra, and the ResearchIT at Iowa State University for migrating the python codes to the web-based application, Scott Nelson at Iowa State for use of the fluorimeter, Daniel Rosenbaum at UT Southwestern for providing the PGS construct, and Neal Alto at UT Southwestern for providing various Arf, Arl, and EspG constructs. Research was supported by funding from the National Institutes of Health (R35-GM128786) and Iowa State University and Roy J. Carver Charitable Trust start-up funds to B.C., Stony Brook University start-up funds to S.C., the Deutsche Forschungsgemeinschaft (DFG), Research Training Group GRK2223, and individual grant RO2414/8-1 to K.R, the National Institutes of Health (R01-DK107733) to D.D.B., and the National Science Foundation (1955260) and National Institutes of Health (R01-GM079383) to J.W.

## Author contributions

B.C. conceived the project and oversaw biochemical work. K.R. oversaw cell biological work. J.W. performed MD simulation and energy calculation. S.Y. purified proteins and performed biochemical experiments. Y.L. and A.B. helped with protein purification and biochemical assays. M.S. performed cellular experiments. B.D. and S.C. helped with structural analysis. D.A.K. performed DLS analysis. L.D., O.A., and D.D.B. helped with cell biological analysis. B.C. wrote the manuscript and prepared figures with assistance from all authors.

## Methods

### Protein purification

All WRC constructs used in this work were derived from previously published WRC230WCA (also called WRC230VCA or WRC^apo^) and ΔWRC230 by standard molecular biology procedures and were verified by Sanger sequencing (Chen et al., 2014a, 2017; Ding et al., 2022). WRC230WCA contains human full-length Sra1, full-length Nap1, WAVE1(1-230)-(GGS)_6_-WCA(485-559), Abi2(1-158), and full-length HSPC300. ΔWRC230 also contains the same subunits except that WAVE1(1-230)-(GGS)_6_-WCA(485-559) is replaced by WAVE1(1-230). Other WRCs contain modified subunits described in detail in **Tables S1** and **S2**.

The WRCs were expressed and purified essentially as previously described (Chen et al., 2014a, 2017). Reconstitution of the recombinant WRC is a multi-step process, involving purification of individual proteins from different host cells (prokaryotic cell and insect cell), assembly/purification of sub-complexes (Sra1/Nap1 dimer and WAVE1/Abi2/HSPC300 trimer) and finally of the WRC pentamer by a series of affinity, ion exchange and gel filtration chromatography steps. Mutations introduced into WRC subunits were carefully chosen and typically made to surface-exposed residues, producing complexes that behaved well and identically to the WT WRC during each step of reconstitution (**Figure S2**). Except Sra1 and Nap1, which were expressed in *Tni* cells using the ESF 921 medium (Expression Systems), other proteins were typically expressed in BL21 (DE3)^T1R^ cells (Sigma) at 18 °C overnight or ArcticExpress^TM^ (DE3) RIL cells (Stratagene) at 10°C for 24 hours. GST-Rac1^QP^ and GST-Rac1^Dead^ were purified by Glutathione Sepharose beads (Cytiva), followed by cation-exchange chromatography through a Source SP15 column and gel filtration through a Hiload Superdex 75 column. GST-Arf1 was purified by Glutathione Sepharose beads, followed by anion-exchange chromatography through a Source Q15 column and gel filtration through a Hiload Superdex 75 column. His8-(GGS)_2_-Arf1 and His6-Tev-EspG were purified by Ni-NTA agarose beads (Qiagen), followed by anion-exchange chromatography through a Source Q15 column and gel filtration through a Hiload Superdex 75 column. Untagged Rac1^QP^ and untagged Rac1^Dead^ were purified by SP Sepharose® Fast Flow beads, followed by a Source SP15 column and a Hiload Superdex 75 gel filtration column. Proteins including Arp2/3 complex, actin, WAVE1 WCA, and TEV protease were purified as previously described (Chen et al., 2014a, 2017; Ismail et al., 2009). All ion exchange and gel filtration chromatography steps were performed using columns from Cytiva on an ÄKTA^TM^ pure protein purification system.

### Non-equilibrium pull-down assay

Non-equilibrium GST pull-down experiments were performed as previously described (Chen et al., 2017). Typically, 100-200 pmol of GST-tagged proteins as baits and 100-200 pmol of WRCs as preys were mixed with 20 µL of Glutathione Sepharose beads (Cytiva) in 1 mL of binding buffer (10 mM HEPES pH 7, 50 mM NaCl, 5% (w/v) glycerol, 0.05% (w/v) Triton X100, 2 mM MgCl_2_, and 5 mM β-mercaptoethanol or 1 mM DTT) at 4 °C for 30 min, followed by three washes using 1 mL of the binding buffer in each time of wash. Bound proteins were eluted with the GST elution buffer (100 mM Tris-HCl pH 8.5, 2 mM MgCl_2_, and 30 mM reduced glutathione) and examined by SDS-PAGE.

### Equilibrium pull-down (EPD) assay

Equilibrium pull-down (EPD) experiments were performed essentially as previously described (Chen et al., 2017). Glutathione Sepharose beads (Cytiva) were first equilibrated in EPD buffer (10 mM HEPES pH 7, 50 mM NaCl, 5% (w/v) glycerol, 2 mM MgCl_2_, and 1 mM DTT) and stored as a 50% (v/v) slurry. Before use, all protein samples were dialyzed against EPD buffer overnight at 4 °C or purified by gel filtration through a column equilibrated with the EPD buffer to maximize buffer match. Each reaction was assembled in 100 µL of total volume of EPD buffer in a 200-µL PCR tube (Axygen), which contained 0.1 µM prey (e.g., WRC), varying concentrations of bait (e.g., GST-Arf1), with or without 100 µM untagged Rac1^QP^ or Rac1^Dead^, 30 µL of the Glutathione Sepharose beads (by aliquoting 60 µL of the 50% (v/v) slurry using a wide-bore pipette tip), and 0.05% (w/v) Triton X100 to facilitate mixing. The reactions were gently mixed at 4 °C on a rotary mixer for 30 min. After a brief centrifugation (∼10,000 g for 10 s) to pellet the beads, 40 µL of the supernatant was immediately transferred to 8 µL of 6 X loading buffer (360 mM Tris-HCl pH 6.8, 12% (w/v) SDS, 60% (w/v) glycerol, 0.012% (w/v) bromophenol blue, and 140 mM freshly added β-mercaptoethanol), and analyzed by Coomassie blue-stained SDS-PAGE gels. The gels were imaged by a ChemiDoc^TM^ XRS + system (BioRAD). Total intensity of the Sra1 and Nap1 bands was quantified by ImageJ (FIJI) to determine unbound WRC. The derived fractional occupancy from 2 to 3 independent experiments was pooled to obtain the binding isotherms for global fitting. The program Prism 8 (GraphPad) was used to fit the binding isotherms using the equation below to obtain dissociation constants 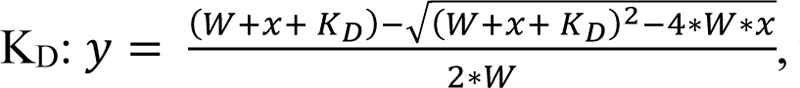 where *y* is the fractional occupancy, *W* is the total WRC concentration (typically 0.1 µM), and x is the total GST-Arf1 concentration.

### Pyrene-actin polymerization assay

Actin polymerization assays were performed as previously described with some modifications here (Chen et al., 2017). Each reaction (120 µL) contained 3-4 µM actin (5% pyrene labeled), 10 nM Arp2/3 complex, 100 nM of various WRC230WCA constructs or WAVE1 WCA, and desired concentrations of untagged Rac1^QP^ and /or His8-Arf1 in the NMEH20GD buffer (50 mM NaCl, 1 mM MgCl_2_, 1 mM EGTA, 10 mM HEPES pH7.0, 20% (w/v) glycerol, and 1 mM DTT). We found that compared to the commonly used KMEI20GD buffer (50 mM KCl, 1 mM MgCl_2_, 1 mM EGTA, 10 mM Imidazole pH7.0, 20% (w/v) glycerol, and 1 mM DTT), the NMEH20GD buffer increased the sensitivity of WRC to Rac1 and Arf1, allowing us to use lower protein concentrations and reduce reaction time in actin assembly assays. Pyrene-actin fluorescence was recorded every 5 seconds at 22 °C, one reaction per measurement using a single-channel pipette to minimize air bubbles or pipetting errors, using a 96-well flat-bottom black plate (Greiner Bio-One™) in a Spark plate reader (Tecan), with excitation at 365 nm and emission at 407 nm (15 nm bandwidth for both wavelengths). Actin assembly rates at the time where the fluorescence intensity is half of the maximum plateau (t_50_) were derived from the kinetic curves using previously published python scripts (Doolittle et al., 2013), which is also implemented on a web application of the scripts (https://biochempy.bb.iastate.edu).

### Dynamic light scattering (DLS) measurement

All experiments were performed on a Wyatt DynaPro NanoStar instrument using Dynamics 7.1.7 software. Sample definitions were as follows: Mw-R model: Globular Proteins; dn/dC (mL/g): 0.185; RG Model: Sphere; Cuvette: Glass Cuvette; Solvent Name, Glycerol 5%. Otherwise, default parameters from the instruments were used, including refractive index and viscosity. Proteins and buffers were filtered using 0.22-µm centrifugal filters right before use to ensure dust was removed from samples. Proteins were mixed directly, and 10 µL were loaded into a quartz microcuvette. Each protein mixture was repeated multiple times, with each repeat undertaking 20 acquisitions (5 seconds per acquisition). The cuvette was cleaned by washing three times with filtered MilliQ water, three times with filtered 95% ethanol, then dried using filtered air. Cutoffs for acceptable runs were defined as any run with SOS (sum of squares) less than 10.0 and with a baseline reading between 0.995 and 1.005. Acquisitions exceeding these values were excluded. For each protein mixture, the readings of all acquisitions from multiple repeats were pooled to obtain the average molecule radius and compared for statistical significance of differences using the ANOVA test in the software R.

### Molecular dynamics simulations

We applied molecular dynamics (MD) simulations and free energies to optimize Arf1 and WRC binding poses. In total, we simulated 6 binding poses that placed the Arf1 close to the D site. Each MD system consists of one WRC bound to two Rac1 molecules (PDB: 7USE) (Ding et al., 2022), one Arf1 (PDB: 1J2J), 400 Cl^-^ and a certain Na^+^ neutralized the systems, and 231710 water molecules. The proteins and cofactors were described by AMBER FF14SB (Maier et al., 2015) and GAFF (Wang et al., 2004) force fields, respectively. MD simulations were performed using a well-established protocol described elsewhere (Kim et al., 2021; Su et al., 2019; Zhang et al., 2021). Briefly, each MD system was first relaxed by a series of minimizations followed by four phases of MD simulations, including the relaxation phase (in total 5 nanoseconds [ns] with 1 femtosecond [fs] time steps), the system heating-up phase (in total 10 ns), the equilibrium phase (10 ns), and the final sampling phase (100 ns). The time step was 2 fs for the last three phases, and the MD simulations of the last two phases were performed at 298K and 1 bar to produce isothermal-isobaric ensembles. All MD simulations were performed using the pmemd.cuda program in AMBER 18 (Case et al., 2018). Besides the root-mean-square deviation (r.m.s.d) ∼ time curves, a representative MD conformation which has the smallest r.m.s.d. between itself and the average MD structure was identified for each MD system.

### Free energy calculations

140 snapshots from the sampling phase (30 – 100 ns) of a trajectory were collected for free energy calculations. An internal program was applied to calculate the MM-PBSA-WSAS free energies of the complex and the binding free energy between Arf1 and WRC. The polar part of the solvation free energy was calculated using Delphi 95 software (Li et al., 2012; Rocchia et al., 2001), and the nonpolar part was estimated by scaling the solvent accessible surface area as described elsewhere (Wang et al., 2019, 2006). The conformational entropy term was predicted using WSAS, a weighted solvent-accessible surface area method (Wang and Hou, 2012). The interior and exterior dielectric constants of PBSA calculations were set to 1.0 and 80.0, respectively. To study the effect of M-site mutations, we conducted computational mutagenesis using the wildtype snapshots and calculated the MM-PBSA-WSAS free energies of complex and Arf1 binding.

### Cell culture and co-immunoprecipitation

B16-F1-derived *Sra1/Cyfip2* KO cells (clone #3) were previously described (Schaks et al., 2018), and maintained in DMEM (4.5 g/l glucose; Invitrogen) supplemented with 10% FCS (Gibco), 2 mM glutamine (Thermo Fisher Scientific) and penicillin (50 Units/ml)/streptomycin (50 µg/ml) (Thermo Fisher Scientific). Cells were routinely transfected in 6 well plates (Sarstedt), using 1 µg DNA in total and 2 µl JetPrime per well.

pEGFP-C2-Sra-1 (CYFIP1) and the derived Y967A mutant construct were described previously (Schaks et al., 2018), and corresponded to the splice variant *CYFIP1a*, sequence AJ567911, of murine origin. Further point mutations in the M site were introduced by site-directed mutagenesis. The identity of all DNA constructs was verified by sequencing.

For EGFP-immunoprecipitation experiments, B16-F1-derived cell lines ectopically expressing EGFP-tagged variants of CYFIP1 were lysed with lysis buffer (1% Triton X-100, 140 mM KCl, 50 mM Tris/HCl pH 7.4 supplemented with 50 mM NaF, 10 mM Na_4_P_2_O_7_, 2 mM MgCl_2_ and Complete Mini, EDTA-free protease inhibitor [Roche]). Lysates were cleared and incubated with GFP-Trap agarose beads (Chromotek) for 60 min. Subsequently, beads were washed three times with lysis buffer lacking protease inhibitor and Triton X-100, mixed with SDS-PAGE loading buffer, boiled for 5 min, and examined by Western Blotting using primary antibodies against Sra-1/Cyfip2 (Steffen et al., 2004), Nap1 (Steffen et al., 2004), WAVE (Schaks et al., 2018) and Abi1 (D3G6C, #39444 Cell Signaling Technology), as well as corresponding, HRP-conjugated secondary antibodies (Invitrogen). Chemiluminescence signals were obtained upon incubation with ECL™ Prime Western Blotting Detection Reagent (Cytiva), and recorded with ECL Chemocam imager (Intas, Goettingen, Germany).

### Fluorescence microscopy, phalloidin staining, and quantifications

B16-F1-derived cell lines expressing indicated, EGFP-tagged CYFIP1 constructs or untransfected control cells were seeded onto laminin-coated (25 µg/ml), 15 mm-diameter glass coverslips and allowed to adhere for about 24 hours prior to fixation. Cells were fixed with pre-warmed, 4% paraformaldehyde (PFA) in PBS for 20 min, and permeabilized with 0.05% Triton-X100 in PBS for 30 sec. The actin cytoskeleton was subsequently stained using ATTO-594-conjugated phalloidin (ATTO TEC GmbH, Germany). Samples were mounted using VectaShield Vibrance antifade reagent and imaged using a 63×/1.4NA Plan apochromatic oil objective.

For assessment of lamellipodia formation, cells were randomly selected and categorized in a blinded manner as follows: “no lamellipodia” if no phalloidin-stained peripheral lamellipodia-like actin meshwork was visible, “immature lamellipodia” if the lamellipodia-like actin meshwork was small, narrow, or displayed multiple ruffles, and “lamellipodia” if the protrusive actin meshworks appeared to be fully developed (see representative examples in Figure 6A).

### Statistical analysis

To assess statistical significance, one-way ANOVA with Dunnett’s post-hoc test was applied to compare multiple groups with one control group. Statistical analyses were performed using Prism 6.01. An error probability below 5% (p < 0.05; * in Figure panels) was considered to imply statistical significance. ** and *** indicated p-values ≤ 0.01 and ≤ 0.001, respectively.

## Supplemental information

**Table S1.**
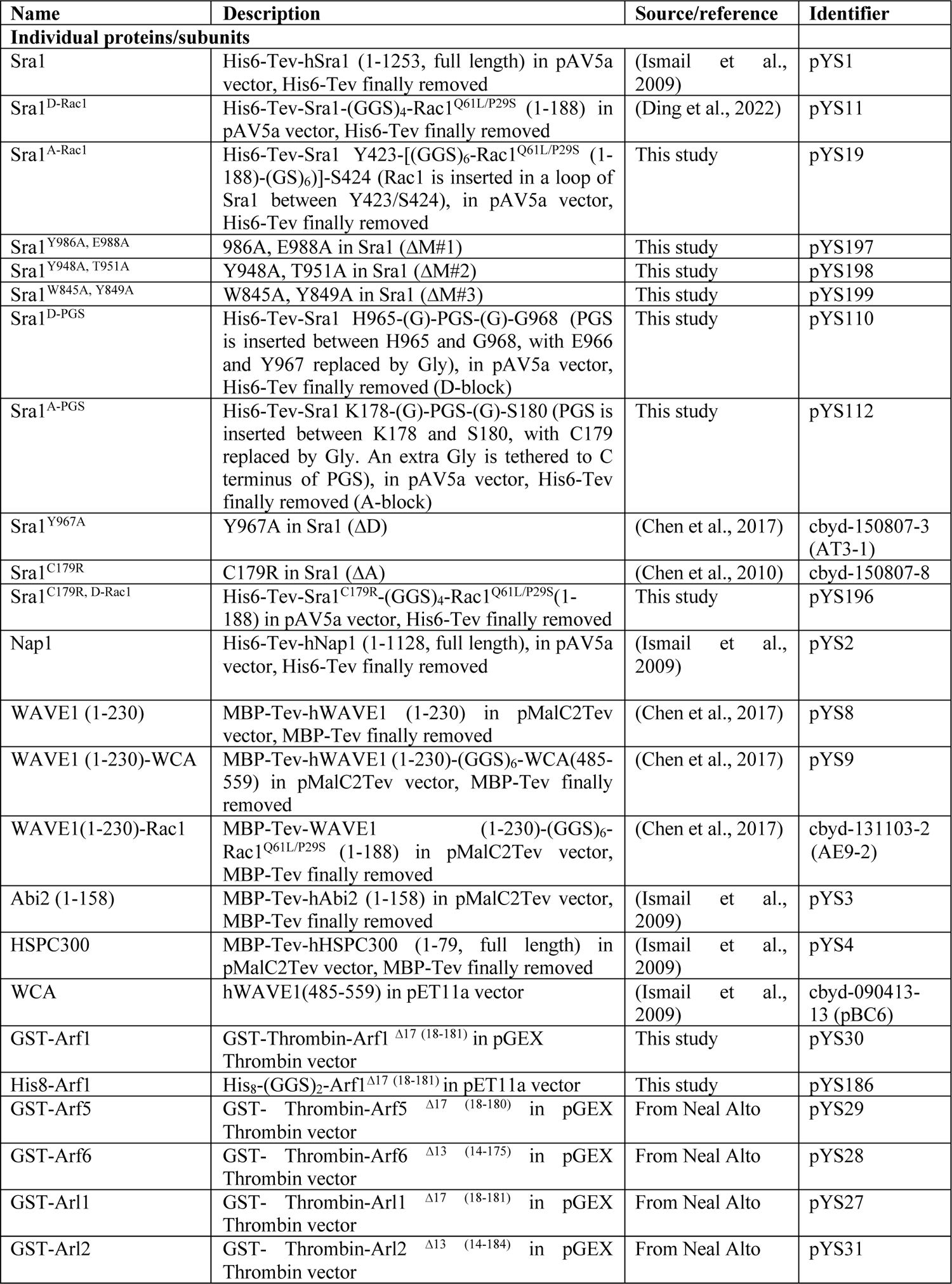
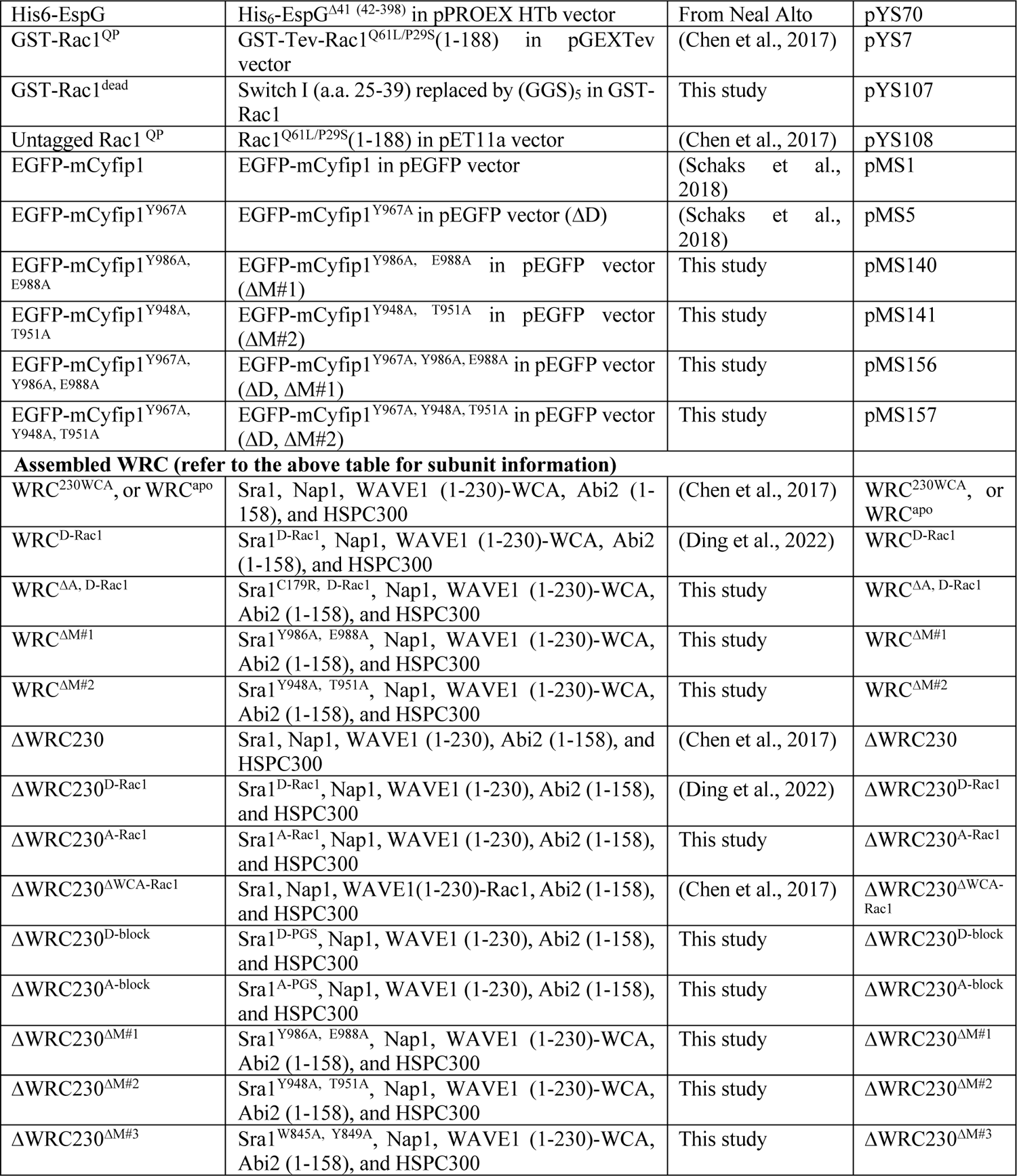
DNA constructs and WRC assemblies used in this study.

**Table S2.**
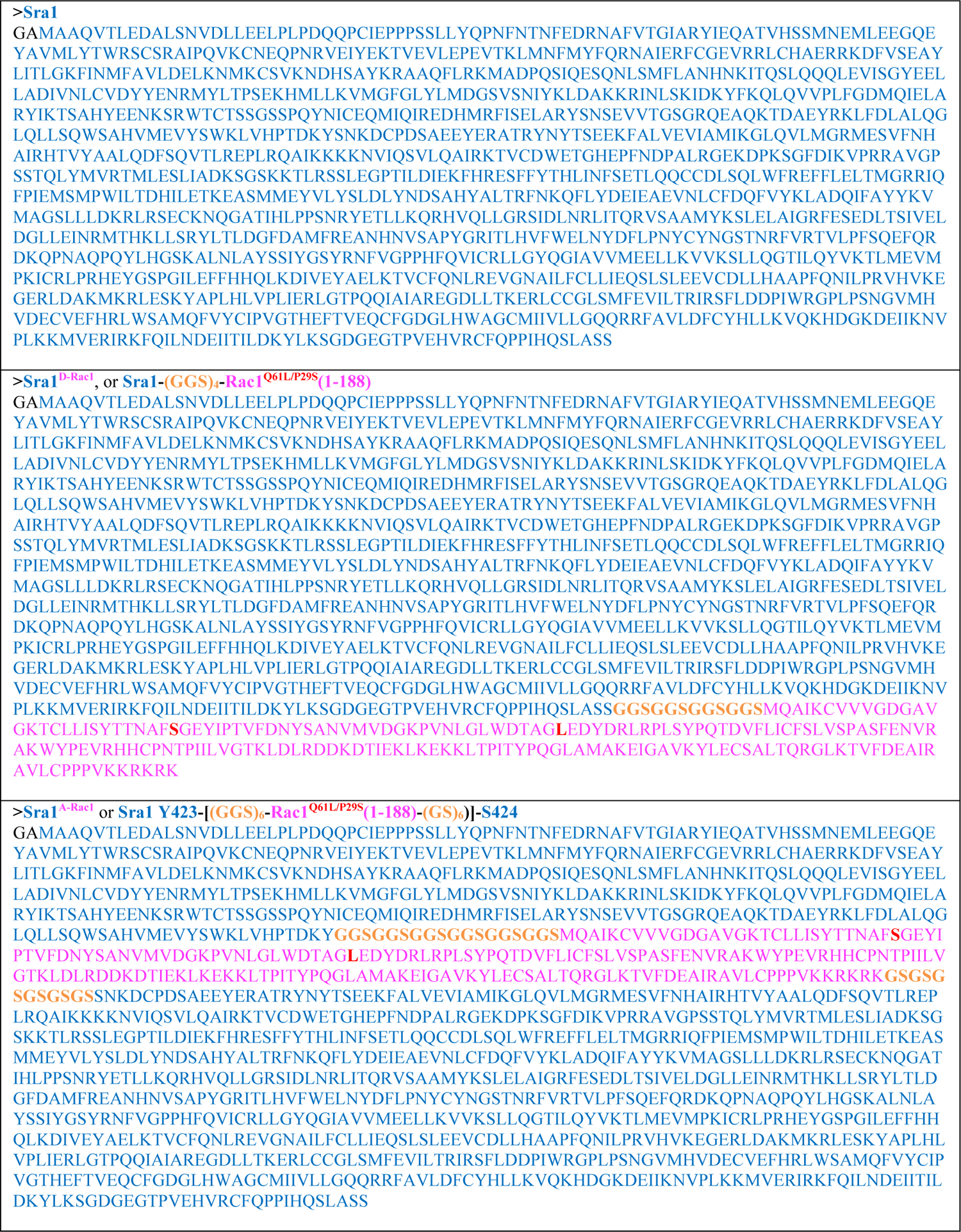
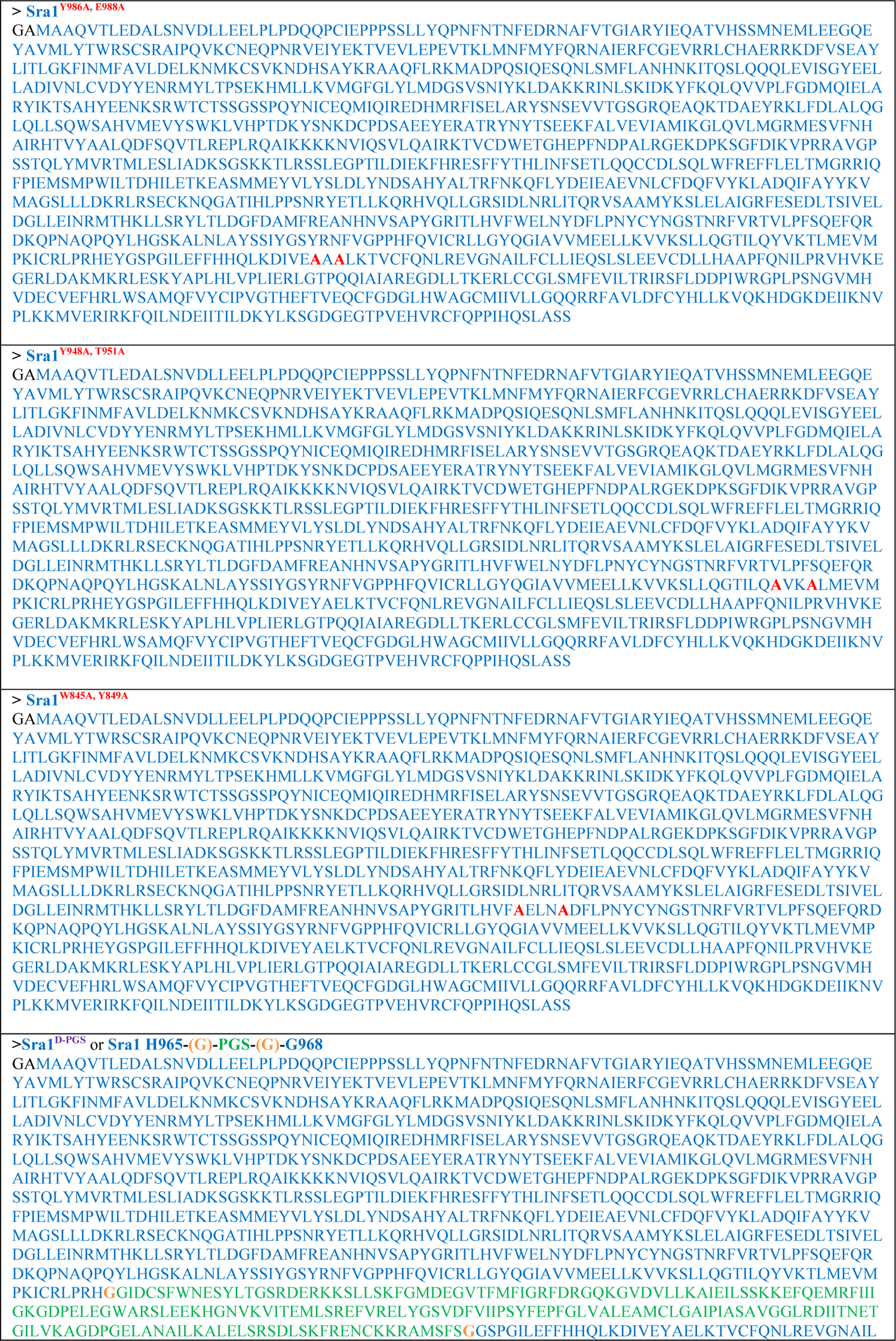
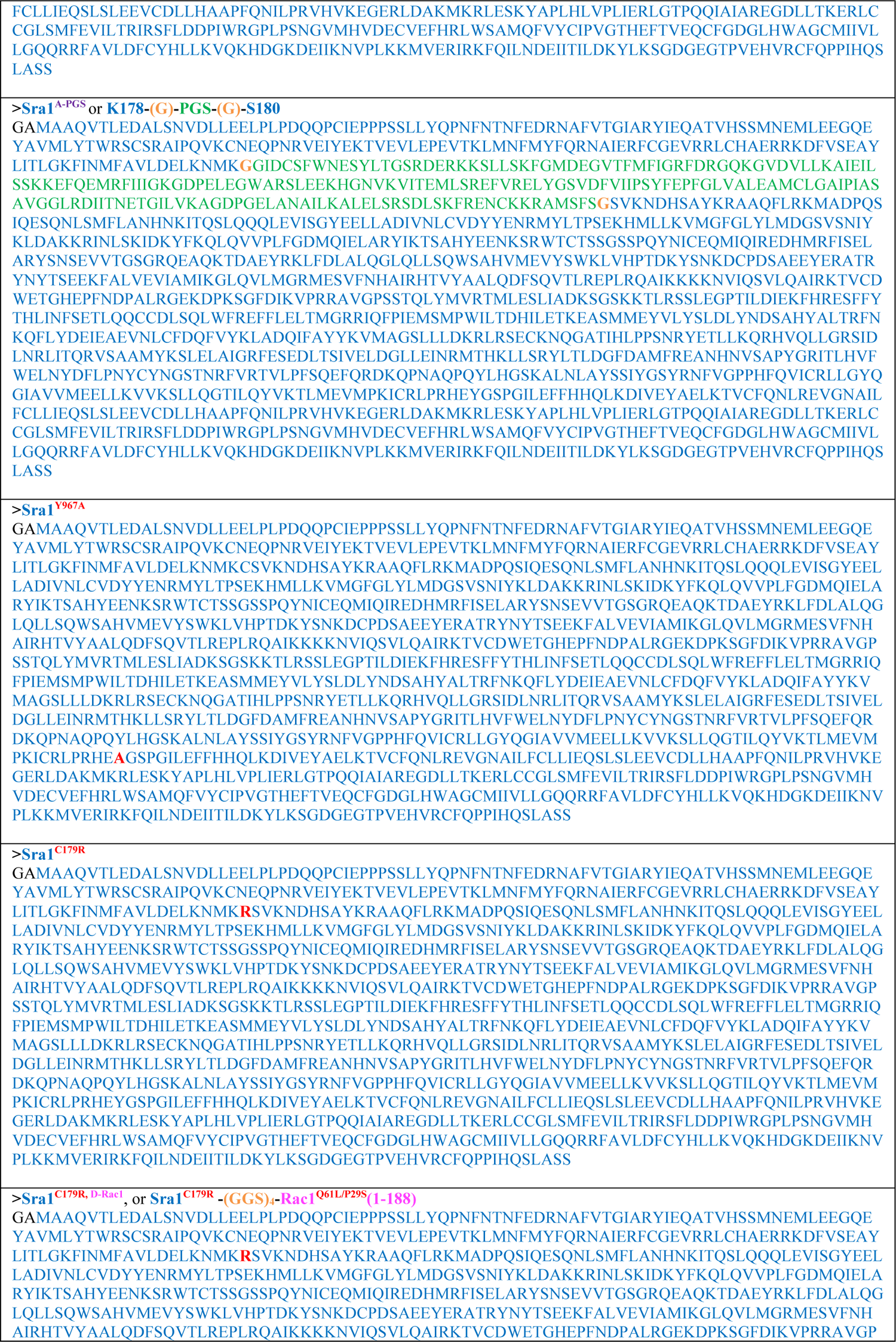
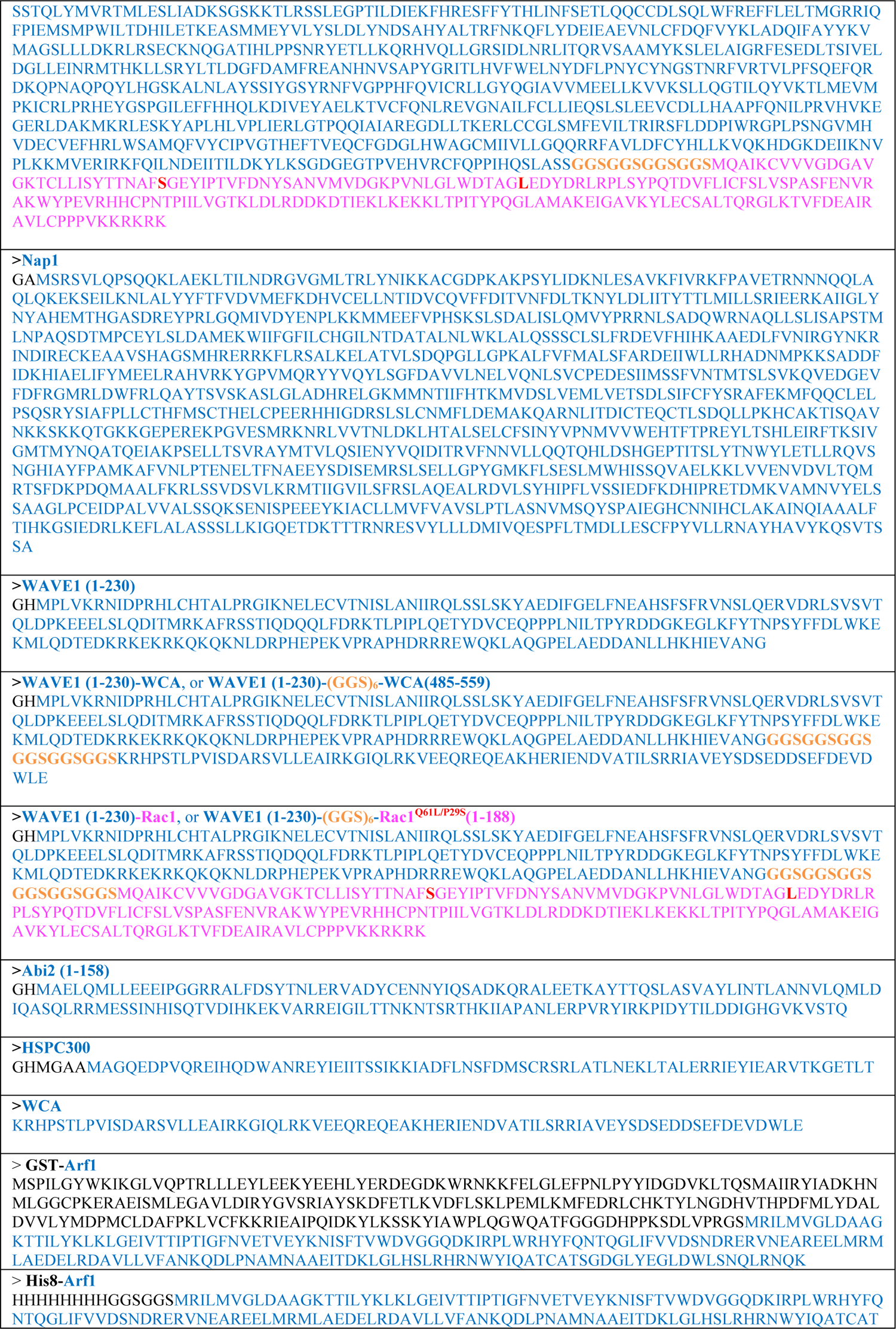
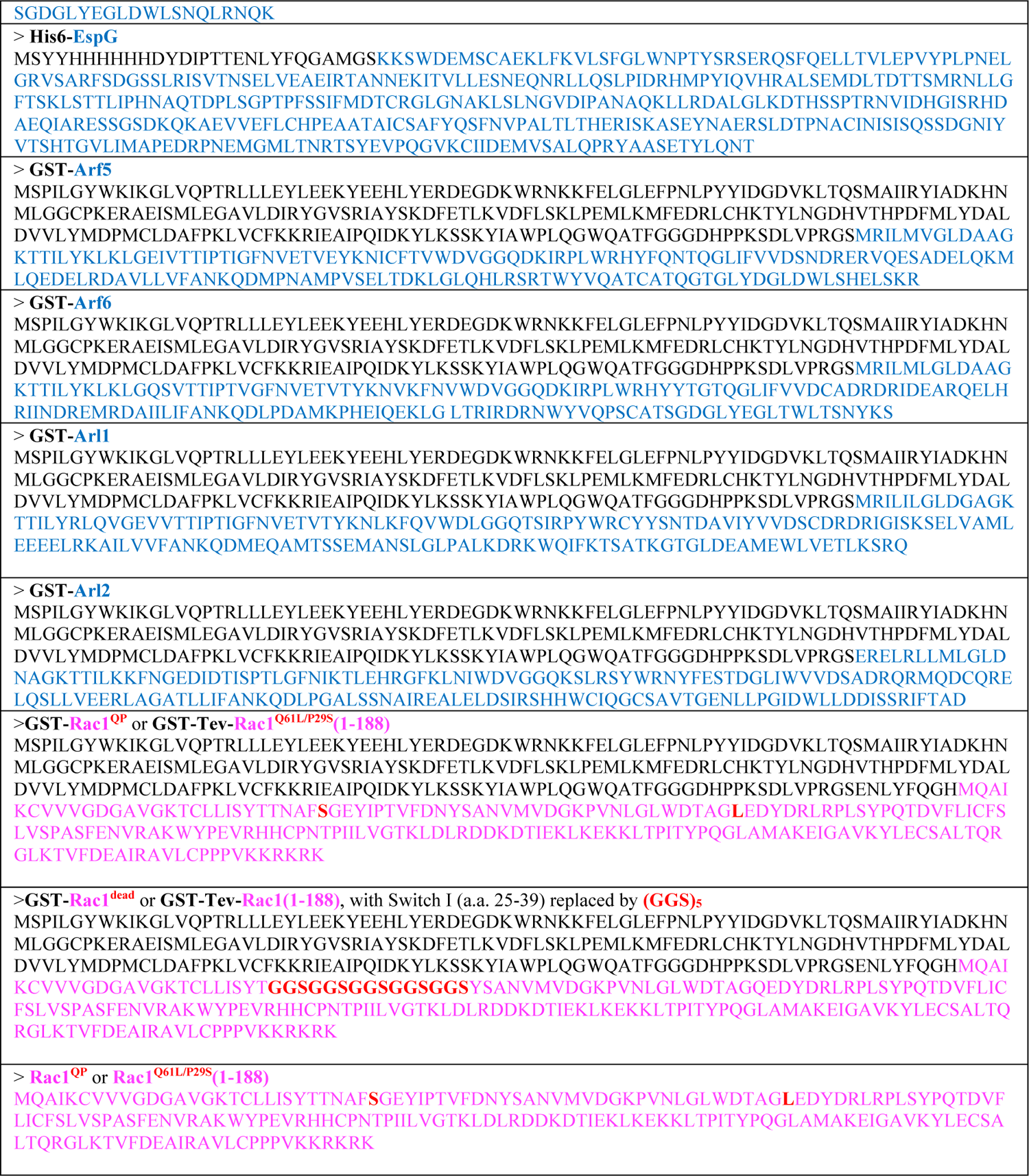
Sequences of recombinant proteins used in this study. Only sequences in the final product (i.e., after protease cleavage to remove the affinity tag) are shown and are annotated by corresponding colors.

---

**Figure S1.**
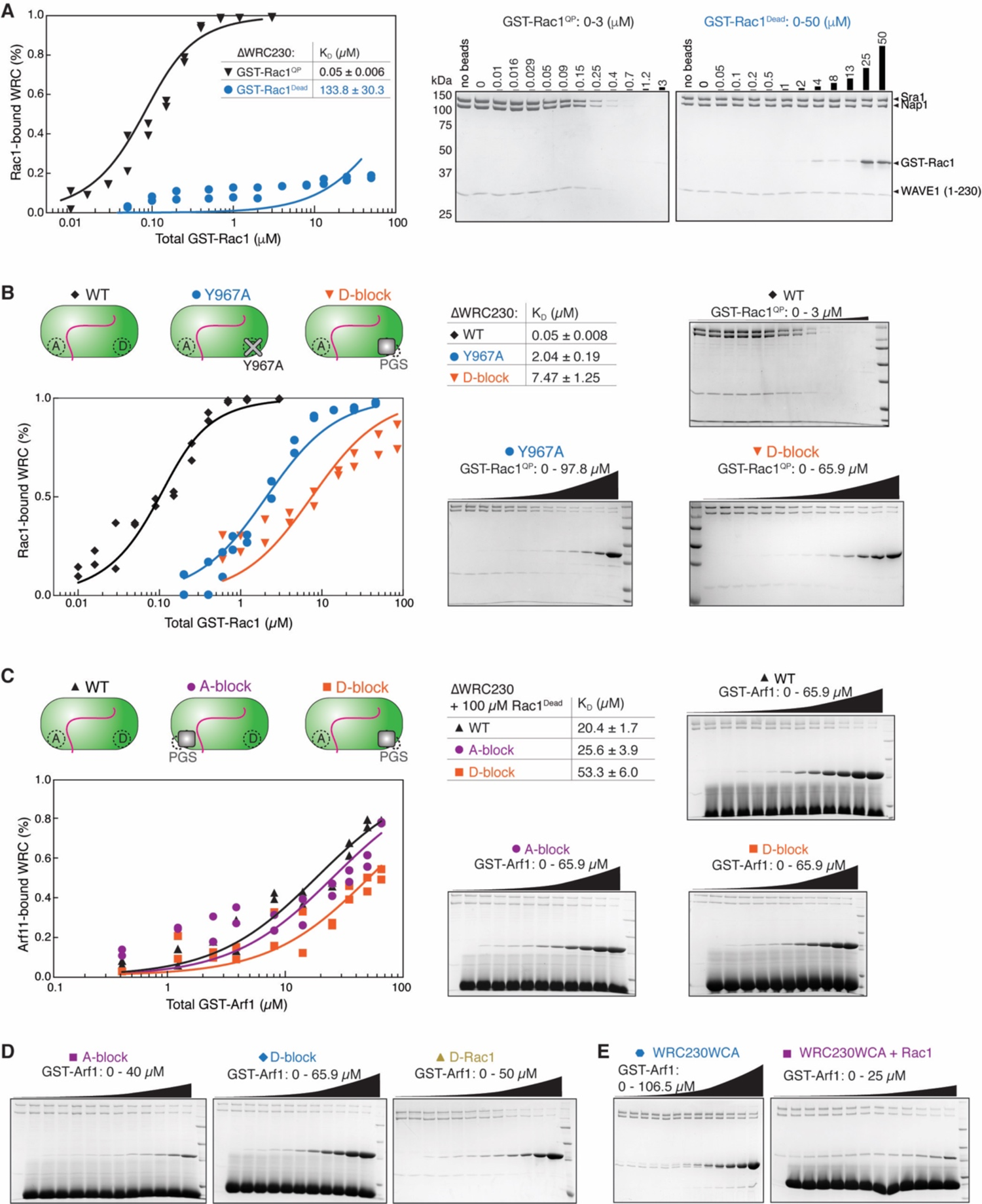
Supporting EDP data for Figures 1, 2, and 3 (Related to Figures 1, 2, and 3). **(A)** EPD measurement, including quantification and representative Coomassie blue-stained SDS PAGE gels, showing GST-Rac1^Dead^ does bind to WRC. **(B)** EPD measurement showing that, compared to the Y967A mutation, D-block further reduces potential leaky binding of Rac1 to the D site. **(C)** EPD measurement showing that blocking the A site or the D site does not significantly affect the basal level binding of Arf1 to WRC. **(D-E)** Example EPD gels for Figures 2B and 3B, respectively.

**Figure S2.**
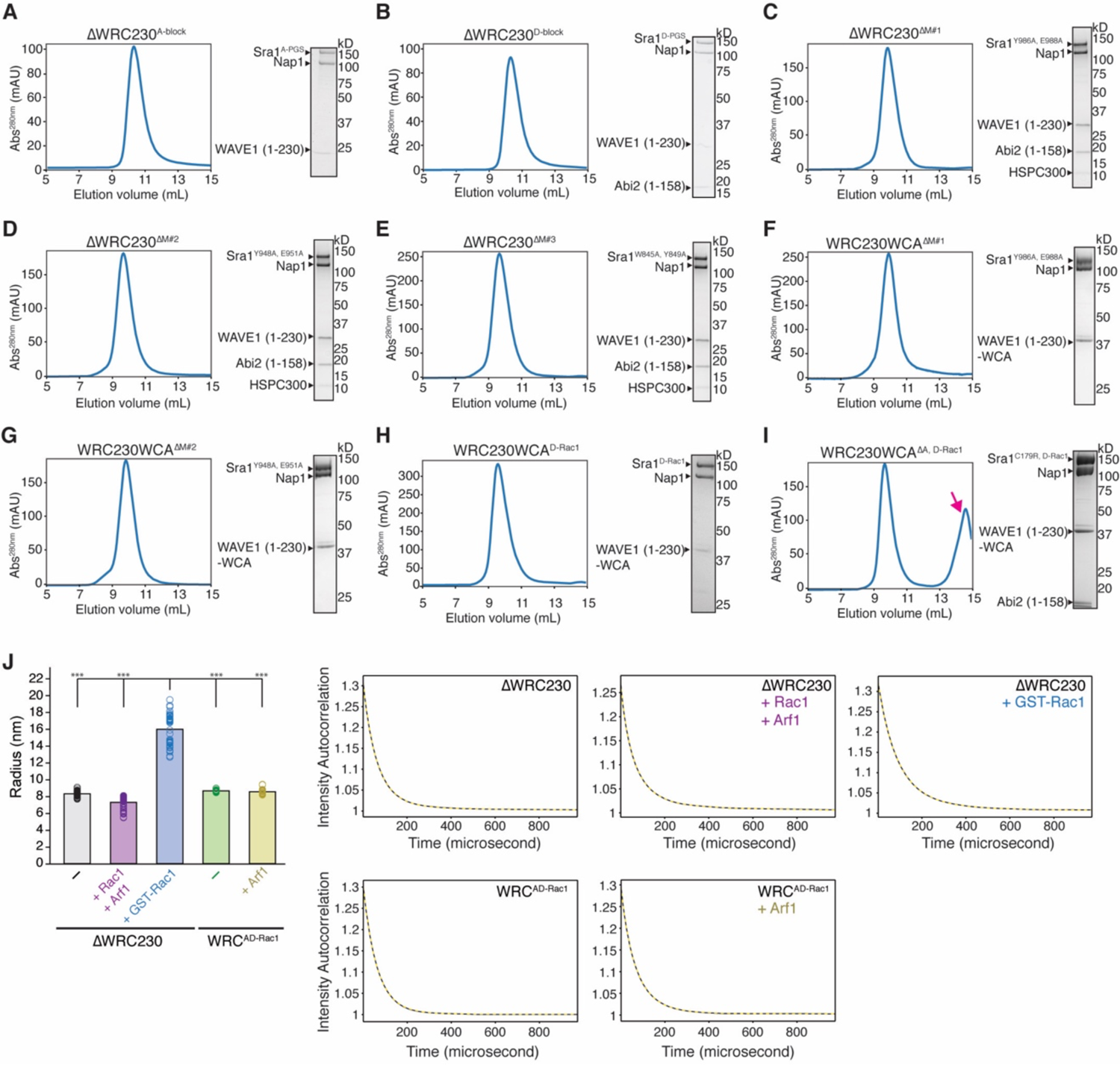
Quality control of the WRCs newly developed in this study (Related to Figures 1, 2, 3, and 5). **(A-I)** Shown are the final steps or analytical steps of WRC purifications using 24-ml Superdex 200 gel filtration columns, with Coomassie blue-stained SDS-PAGE gels showing the peak or pooled fractions. Depending on whether the preceding purification step included a Source Q15 ion exchange column, different amounts of Tev and cleaved MBP tag may show as clearly separated peaks (indicated by magenta arrow) following the WRC peak. **(J)** Dynamic Light Scattering (DLS) measurements of indicated WRC mixed with GTPases. GST-Rac1 is used as a positive control to show radius change when WRC is dimerized by GST-Rac1. On the left is radius values for 3.3 µM ΔWRC230 alone (n = 54), 3.5 µM ΔWRC230 + 30 µM Rac1^QP^ + 125 µM Arf1 (n = 44), 1 µM ΔWRC230 + 30 µM GST-Rac1^QP^ (n = 51), 5 µM WRC^AD-Rac1^ alone (n = 16), and 5 µM WRC^AD-Rac1^ + 125 µM Rac1^QP^ (n = 24). n equals total number of acquisitions, *** indicates p < 0.001, ANOVA test. Radius values are reported as the average values from all requisitions. All experiments were collected in 50-100 mM NaCl and 5% (w/v) glycerol and at room temperature. The slightly reduced radius for ΔWRC230 + Rac1 + Arf1 sample was likely due to the addition of large amounts of Rac1 and Arf1, which have lower molecular weight. On the right is a representative plot of intensity autocorrelation (black solid curve) and the regularization fit (yellow dashed curve) for each experiment.

**Figure S3.**
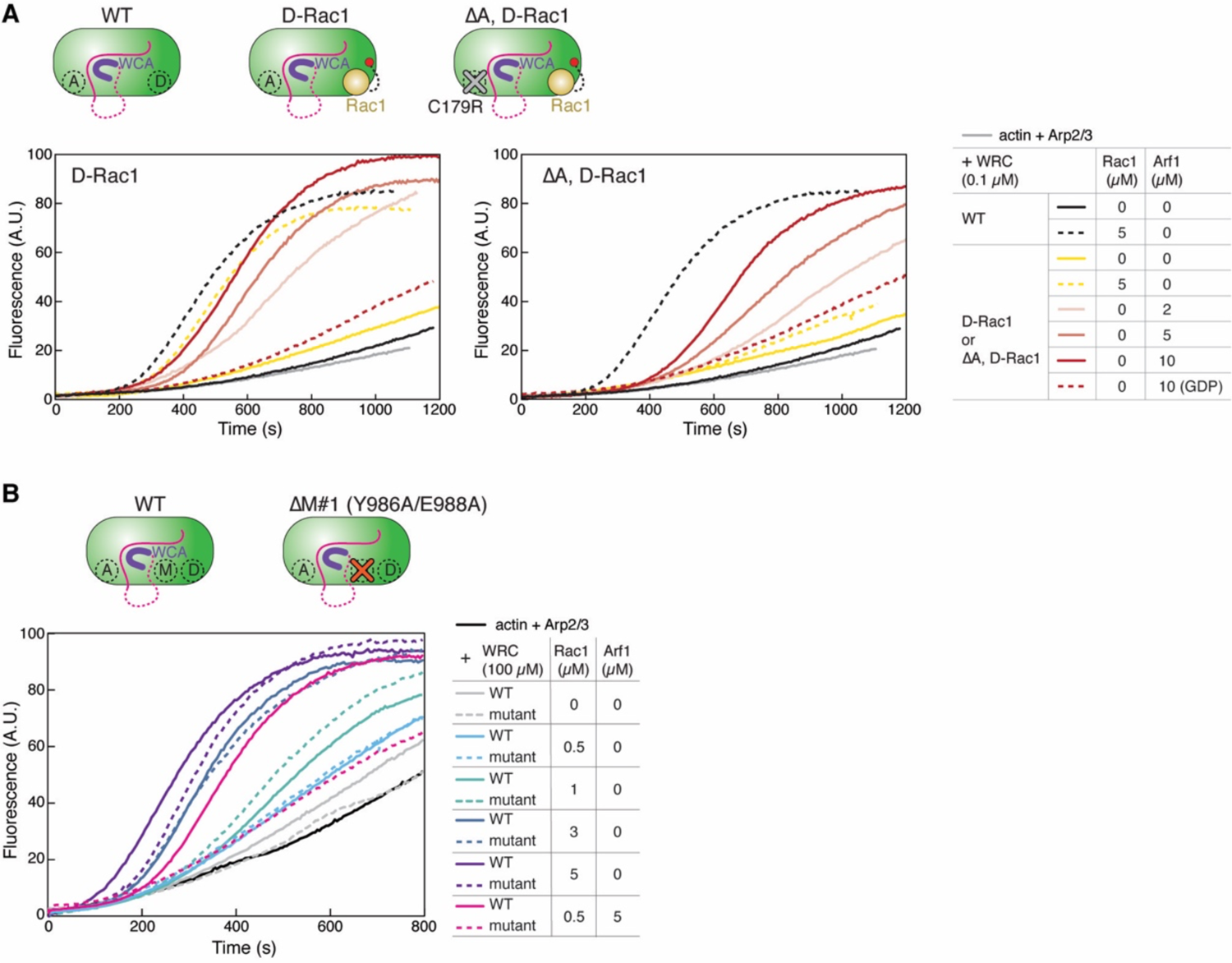
Additional actin polymerization assays comparing Arf1-vs. Rac1-mediated WRC activation (Related to Figures 3 and 5). **(A-B)** Pyrene-actin polymerization assays comparing indicated WRC variants in response to the addition of free Rac1^QP^ or Arf1. WT WRC230WCA was used as a reference point. Reactions contained 3.5 µM actin (5% pyrene labeled), 10 nM Arp2/3 complex, 100 nM indicated WRC, and indicated amounts of Rac1^QP^ and/or Arf1. In all assays, Arf1 is loaded with GNPPNP, unless it is designated with GDP.

**Figure S4.**
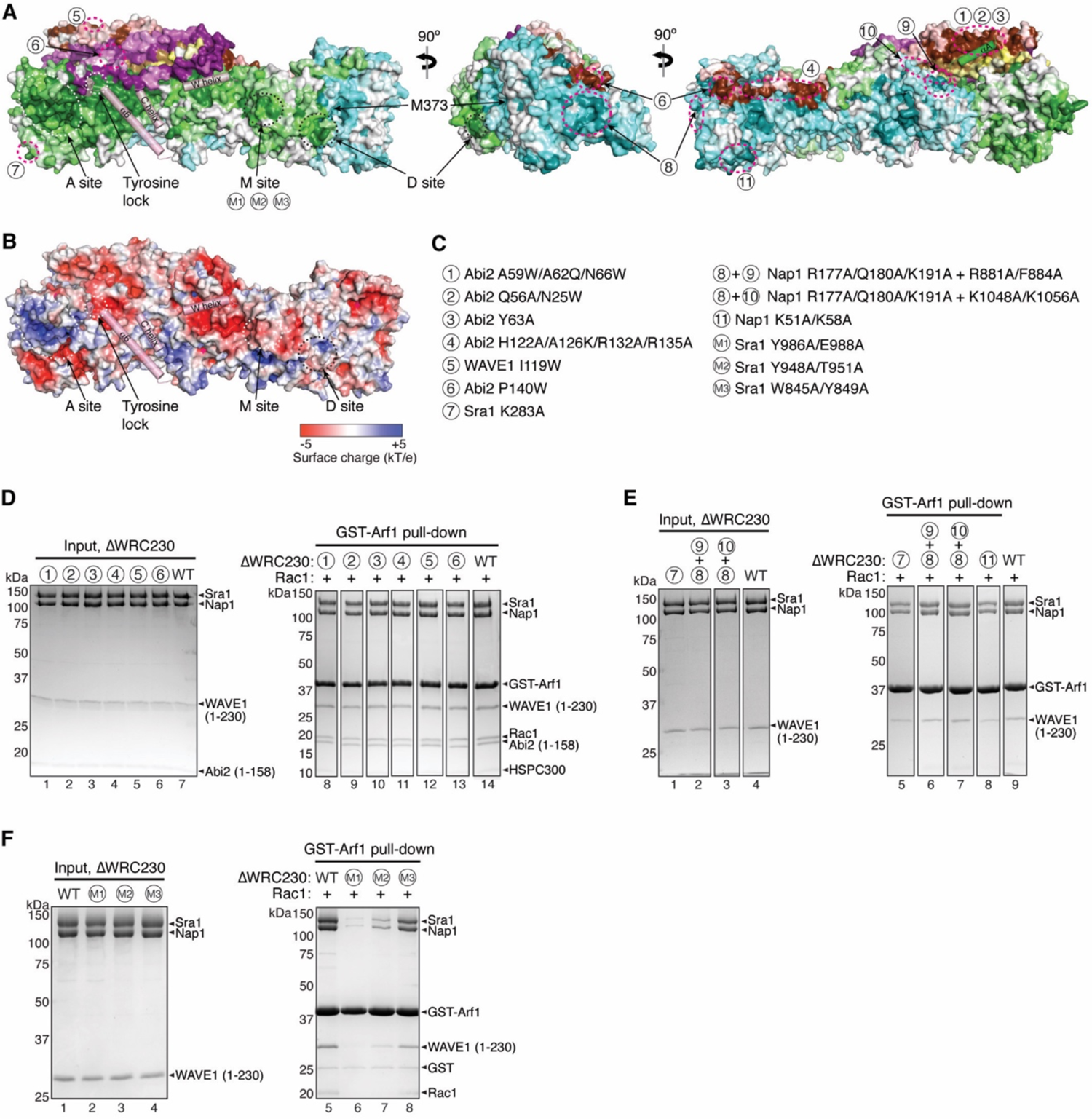
Surveying various conserved surfaces on WRC to identify the Arf1-binding site (Related to Figures 4 and 5). **(A-B)** Surface conservation (A) and electrostatic charge (B) representation of the WRC. (A) was calculated by Consurf (Ashkenazy et al., 2016), and (B) by APBS in Pymol (Jurrus et al., 2018). In (A), color to white gradients represent the most conserved surface residues (ConSurf score = 9 for darkest colors) to the least conserved residues (ConSurf score = 1 for white color). Important sites on Sra1, including the identified Arf1 binding site (M site), are indicated with white or black dotted circles. Positions of mutations examined in (D-F) are indicated with magenta dotted circles. Semitransparent pink cylinders refer to the sequences in WAVE1 that are destabilized upon WRC activation by Rac1. The *α*A helix in Sra1 is also shown as a semitransparent green cylinder for clarity. **(C)** Information of the mutations indicated in (A) and examined in (D-F). **(D-F)** Coomassie blue-stained SDS PAGE gels showing GST-Arf1 pull-down of ΔWRC230 carrying indicated surface mutations.

**Figure S5.**
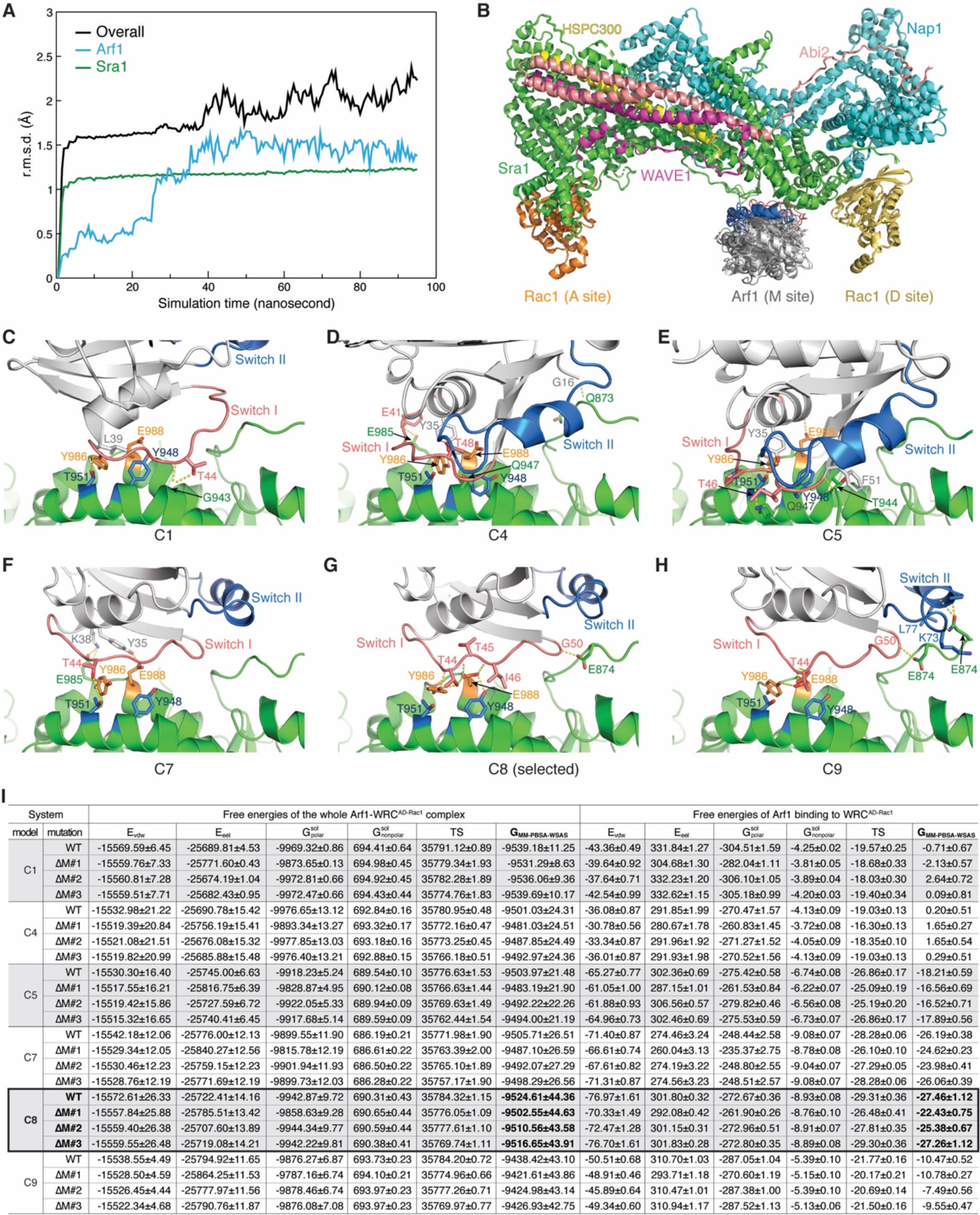
Molecular dynamics simulation and energy minimization of the top M-site Arf1 docking models (Related to Figure 4). **(A)** Root-mean-square deviation (r.m.s.d.) time course of the MD simulation of the C8 model, showing the system reached equilibrium after 30 nanoseconds. Other models showed similar r.m.s.d. time courses. **(B)** Overlay of all six MD energy minimized models. **(C-H)** Detailed view of the Arf1-M site interactions in indicated models. Sra1 is kept in the same orientation to demonstrate different orientations of Arf1 in various MD models. Yellow dotted lines indicate polar or *π*-*π* interactions. **(I)** List of MM-PBSA-WSAS energy terms of whole complex free energies (left) and Arf1-binding free energies (right) of six Arf1-WRC^AD-Rac1^ complexes, each including the WT WRC and M-site mutants, with ΔM#1 for Sra1^Y986A/E988A^, ΔM#2 for Sra1^948A/T951A^, and ΔM#3 for Sra1^W845A/Y849A^ (as a negative control). All energy terms, including E_vdw_ and E_eel_ for the van der Waals and electrostatic interactions, respectively; G^sol^_polar_ and G^sol^_nonpolar_ for the polar and nonpolar parts of the solvation free energy, respectively, and TS for the entropic term, are in kcal/mol. Model C8 agreed best with the experimental results, in that ΔM#1 and ΔM#2, but not ΔM#3 mutations, increased by free energies for Arf1 binding.

## References

1. Anitei, M., Stange, C., Parshina, I., Baust, T., Schenck, A., Raposo, G., Kirchhausen, T., and Hoflack, B. (2010). Protein complexes containing CYFIP/Sra/PIR121 coordinate Arf1 and Rac1 signalling during clathrin-AP-1-coated carrier biogenesis at the TGN. Nat. Cell Biol. 12.

2. Ashkenazy, H., Abadi, S., Martz, E., Chay, O., Mayrose, I., Pupko, T., and Ben-Tal, N. (2016). ConSurf 2016: an improved methodology to estimate and visualize evolutionary conservation in macromolecules. Nucleic Acids Res. 44.

3. Balasubramanian, N., Scott, D.W., Castle, J.D., Casanova, J.E., and Schwartz, M.A. (2007). Arf6 and microtubules in adhesion-dependent trafficking of lipid rafts. Nat. Cell Biol. 9.

4. Baust, T., Czupalla, C., Krause, E., Bourel-Bonnet, L., and Hoflack, B. (2006). Proteomic analysis of adaptor protein 1A coats selectively assembled on liposomes. Proc. Natl. Acad. Sci. U. S. A. 103.

5. Boshans, R.L., Szanto, S., van Aelst, L., and D’Souza-Schorey, C. (2000). ADP-Ribosylation Factor 6 Regulates Actin Cytoskeleton Remodeling in Coordination with Rac1 and RhoA. Mol. Cell. Biol. 20.

6. Case, D.A., Walker, R.C., Cheatham, T.E., Simmerling, C., Roitberg, A., Merz, K.M., Luo, R., and Darden, T. (2018). Amber 2018. Univ. California, San Fr. 2018.

7. Chen, B., Padrick, S.B., Henry, L., and Rosen, M.K. (2014a). Biochemical reconstitution of the WAVE regulatory complex. Methods Enzymol. 540, 55–72.

8. Chen, B., Brinkmann, K., Chen, Z., Pak, C.W., Liao, Y., Shi, S., Henry, L., Grishin, N. V., Bogdan, S., and Rosen, M.K. (2014b). The WAVE regulatory complex links diverse receptors to the actin cytoskeleton. Cell 156, 195–207.

9. Chen, B., Chou, H.T., Brautigam, C.A., Xing, W., Yang, S., Henry, L., Doolittle, L.K., Walz, T., and Rosen, M.K. (2017). Rac1 GTPase activates the WAVE regulatory complex through two distinct binding sites. Elife 6.

10. Chen, E.H., Pryce, B.A., Tzeng, J.A., Gonzalez, G.A., and Olson, E.N. (2003). Control of myoblast fusion by a guanine nucleotide exchange factor, loner, and its effector ARF6. Cell 114.

11. Chen, X.J., Squarr, A.J., Stephan, R., Chen, B., Higgins, T.E., Barry, D.J., Martin, M.C., Rosen, M.K., Bogdan, S., and Way, M. (2014c). Ena/VASP proteins cooperate with the WAVE complex to regulate the actin cytoskeleton. Dev. Cell 30, 569–584.

12. Chen, Z., Borek, D., Padrick, S.B., Gomez, T.S., Metlagel, Z., Ismail, A.M., Umetani, J., Billadeau, D.D., Otwinowski, Z., and Rosen, M.K. (2010). Structure and Control of the Actin Regulatory WAVE Complex. Nature 468, 533–538.

13. Cheng, A., Arumugam, T. V., Liu, D., Khatri, R.G., Mustafa, K., Kwak, S., Ling, H.P., Gonzales, C., Xin, O., Jo, D.G., et al. (2007). Pancortin-2 interacts with WAVE1 and Bcl-xL in a mitochondria-associated protein complex that mediates ischemic neuronal death. J. Neurosci. 27.

14. Cook, S.A., Comrie, W.A., Poli, M.C., Similuk, M., Oler, A.J., Faruqi, A.J., Kuhns, D.B., Yang, S., Vargas-Hernández, A., Carisey, A.F., et al. (2020). HEM1 deficiency disrupts mTORC2 and F-actin control in inherited immunodysregulatory disease. Science (80-.). 369, 202–207.

15. D’Souza-Schorey, C., and Chavrier, P. (2006). ARF proteins: roles in membrane traffic and beyond. Nat Rev Mol Cell Biol 7, 347–358.

16. D’Souza-Schorey, C., Boshans, R.L., McDonough, M., Stahl, P.D., and Van Aelst, L. (1997). A role for POR1, a Rac1-interacting protein, in ARF6-mediated cytoskeletal rearrangements. EMBO J. 16.

17. Derivery, E., Lombard, B., Loew, D., and Gautreau, A. (2009). The Wave complex is intrinsically inactive. Cell Motil. Cytoskelet. 66, 777–790.

18. Desta, I.T., Porter, K.A., Xia, B., Kozakov, D., and Vajda, S. (2020). Performance and Its Limits in Rigid Body Protein-Protein Docking. Structure 28.

19. Ding, B., Yang, S., Schaks, M., Liu, Y., Brown, A., Rottner, K., Chowdhury, S., and Chen, B. (2022). Structures reveal a key mechanism of WAVE Regulatory Complex activation by Rac1 GTPase. BioRxiv *[Preprint]*, May 10. doi: https://doi.org/10.1101/2022.05.10.49.

20. Donaldson, J.G., and Jackson, C.L. (2011). ARF family G proteins and their regulators: Roles in membrane transport, development and disease. Nat. Rev. Mol. Cell Biol.

21. Dong, N., Zhu, Y., Lu, Q., Hu, L., Zheng, Y., and Shao, F. (2012). Structurally distinct bacterial TBC-like GAPs link Arf GTPase to Rab1 inactivation to counteract host defenses. Cell.

22. Doolittle, L.K., Rosen, M.K., and Padrick, S.B. (2013). Measurement and analysis of in vitro actin polymerization. Methods Mol Biol 1046, 273–293.

23. Eden, S., Rohatgi, R., Podtelejnikov, A. V., Mann, M., and Kirschner, M.W. (2002). Mechanism of regulation of WAVE1-induced actin nucleation by Rac1 and Nck. Nature 418, 790–793.

24. Etienne-Manneville, S., and Hall, A. (2002). Rho GTPases in cell biology. Nature 420.

25. Gautreau, A., Ho, H.Y.H., Li, J., Steen, H., Gygi, S.P., and Kirschner, M.W. (2004). Purification and architecture of the ubiquitous Wave complex. Proc. Natl. Acad. Sci. U. S. A. 101.

26. Gillingham, A.K., and Munro, S. (2007). The small G proteins of the Arf family and their regulators. Annu. Rev. Cell Dev. Biol. 23.

27. Honda, A., Nogami, M., Yokozeki, T., Yamazaki, M., Nakamura, H., Watanabe, H., Kawamoto, K., Nakayama, K., Morris, A.J., Frohman, M.A., et al. (1999). Phosphatidylinositol 4-phosphate 5-kinase α is a downstream effector of the small G protein ARF6 in membrane ruffle formation. Cell 99.

28. Hu, B., Shi, B., Jarzynka, M.J., Yiin, J.J., D’Souza-Schorey, C., and Cheng, S.Y. (2009). ADP-ribosylation factor 6 regulates glioma cell invasion through the IQ-domain GTPase-activating protein 1-Rac1-mediated pathway. Cancer Res. 69.

29. Humphreys, D., Liu, T., Davidson, A.C., Hume, P.J., and Koronakis, V. (2012a). The Drosophila Arf1 homologue Arf79F is essential for lamellipodium formation. J Cell Sci 125, 5630–5635.

30. Humphreys, D., Davidson, A., Hume, P.J., and Koronakis, V. (2012b). Salmonella virulence effector SopE and Host GEF ARNO cooperate to recruit and activate WAVE to trigger bacterial invasion. Cell Host Microbe 11, 129–139.

31. Humphreys, D., Davidson, A.C., Hume, P.J., Makin, L.E., and Koronakis, V. (2013). Arf6 coordinates actin assembly through the WAVE complex, a mechanism usurped by Salmonella to invade host cells. Proc Natl Acad Sci U S A 110, 16880–16885.

32. Humphreys, D., Singh, V., and Koronakis, V. (2016). Inhibition of WAVE Regulatory Complex Activation by a Bacterial Virulence Effector Counteracts Pathogen Phagocytosis. Cell Rep 17, 697–707.

33. Hunt, E.L., Rai, H., and Harris, T.J.C. (2022). SCAR/WAVE complex recruitment to a supracellular actomyosin cable by myosin activators and a junctional Arf-GEF during Drosophila dorsal closure. Mol. Biol. Cell.

34. Ismail, A.M., Padrick, S.B., Chen, B., Umetani, J., and Rosen, M.K. (2009). The WAVE Regulatory Complex is Inhibited. Nat. Struct. Mol. Biol. 16, 561–563.

35. Jurrus, E., Engel, D., Star, K., Monson, K., Brandi, J., Felberg, L.E., Brookes, D.H., Wilson, L., Chen, J., Liles, K., et al. (2018). Improvements to the APBS biomolecular solvation software suite. Protein Sci. 27.

36. Kang, R., Tang, D., Yu, Y., Wang, Z., Hu, T., Wang, H., and Cao, L. (2010). WAVE1 regulates Bcl-2 localization and phosphorylation in leukemia cells. Leukemia 24.

37. Kim, P., Li, H., Wang, J., and Zhao, Z. (2021). Landscape of drug-resistance mutations in kinase regulatory hotspots. Brief. Bioinform. 22.

38. Koo, T.H., Eipper, B.A., and Donaldson, J.G. (2007). Arf6 recruits the Rac GEF Kalirin to the plasma membrane facilitating Rac activation. BMC Cell Biol. 8.

39. Koronakis, V., Hume, P.J., Humphreys, D., Liu, T., Hørning, O., Jensen, O.N., and McGhie, E.J. (2011). WAVE regulatory complex activation by cooperating GTPases Arf and Rac1. Proc. Natl. Acad. Sci. U. S. A. 108, 14449–14454.

40. Krauss, M., Kinuta, M., Wenk, M.R., De Camilli, P., Takei, K., and Haucke, V. (2003). ARF6 stimulates clathrin/AP-2 recruitment to synaptic membranes by activating phosphatidylinositol phosphate kinase type Iγ. J. Cell Biol. 162.

41. Lebensohn, A.M., and Kirschner, M.W. (2009). Activation of the WAVE complex by coincident signals controls actin assembly. Mol. Cell 36, 512.

42. Lewis-Saravalli, S., Campbell, S., and Claing, A. (2013). ARF1 controls Rac1 signaling to regulate migration of MDA-MB-231 invasive breast cancer cells. Cell Signal 25, 1813–1819.

43. Li, L., Li, C., Sarkar, S., Zhang, J., Witham, S., Zhang, Z., Wang, L., Smith, N., Petukh, M., and Alexov, E. (2012). DelPhi: a comprehensive suite for DelPhi software and associated resources. BMC Biophys. 5.

44. Maier, J.A., Martinez, C., Kasavajhala, K., Wickstrom, L., Hauser, K.E., and Simmerling, C. (2015). ff14SB: Improving the Accuracy of Protein Side Chain and Backbone Parameters from ff99SB. J. Chem. Theory Comput. 11.

45. Marchesin, V., Montagnac, G., and Chavrier, P. (2015). ARF6 promotes the formation of Rac1 and WAVE-dependent ventral F-actin rosettes in breast cancer cells in response to epidermal growth factor. PLoS One 10, e0121747.

46. Mosaddeghzadeh, N., and Ahmadian, M.R. (2021). The rho family gtpases: Mechanisms of regulation and signaling. Cells 10.

47. Myers, K.R., and Casanova, J.E. (2008). Regulation of actin cytoskeleton dynamics by Arf-family GTPases. Trends Cell Biol. 18.

48. Padrick, S.B., Cheng, H.C., Ismail, A.M., Panchal, S.C., Doolittle, L.K., Kim, S., Skehan, B.M., Umetani, J., Brautigam, C.A., Leong, J.M., et al. (2008). Hierarchical Regulation of WASP/WAVE Proteins. Mol. Cell 32, 426–438.

49. Palacios, F., Schweitzer, J.K., Boshans, R.L., and D’Souza-Schorey, C. (2002). ARF6-GTP recruits Nm23-H1 to facilitate dynamin-mediated endocytosis during adherens junctions disassembly. Nat. Cell Biol. 4.

50. Phuyal, S., and Farhan, H. (2019). Multifaceted Rho GTPase Signaling at the Endomembranes. Front. Cell Dev. Biol. 7.

51. Pollard, T.D. (2010). A guide to simple and informative binding assays. Mol Biol Cell 21, 4061–4067.

52. Quignot, C., Rey, J., Yu, J., Tufféry, P., Guerois, R., and Andreani, J. (2018). InterEvDock2: An expanded server for protein docking using evolutionary and biological information from homology models and multimeric inputs. Nucleic Acids Res. 46.

53. Radhakrishna, H., Al-Awar, O., Khachikian, Z., and Donaldson, J.G. (1999). ARF6 requirement for Rac ruffling suggests a role for membrane trafficking in cortical actin rearrangements. J. Cell Sci. 112.

54. Ramírez-Aportela, E., López-Blanco, J.R., and Chacón, P. (2016). FRODOCK 2.0: Fast protein-protein docking server. Bioinformatics 32.

55. Ren, X., Farías, G.G., Canagarajah, B.J., Bonifacino, J.S., and Hurley, J.H. (2013). Structural basis for recruitment and activation of the AP-1 clathrin adaptor complex by Arf1. Cell 152.

56. Rocchia, W., Alexov, E., and Honig, B. (2001). Extending the applicability of the nonlinear Poisson-Boltzmann equation: Multiple dielectric constants and multivalent ions. J. Phys. Chem. B 105.

57. Rottner, K., Stradal, T.E.B., and Chen, B. (2021). WAVE regulatory complex. Curr. Biol. 31, R512–R517.

58. Santy, L.C., and Casanova, J.E. (2001). Activation of ARF6 by ARNO stimulates epithelial cell migration through downstream activation of both Rac1 and phospholipase D. J. Cell Biol. 154.

59. Santy, L.C., Ravichandran, K.S., and Casanova, J.E. (2005). The DOCK180/Elmo complex couples ARNO-mediated Arf6 activation to the downstream activation of Rac1. Curr. Biol. 15.

60. Schaks, M., Singh, S.P., Kage, F., Thomason, P., Klünemann, T., Steffen, A., Blankenfeldt, W., Stradal, T.E., Insall, R.H., and Rottner, K. (2018). Distinct Interaction Sites of Rac GTPase with WAVE Regulatory Complex Have Non-redundant Functions in Vivo. Curr. Biol. 28.

61. Schaks, M., Reinke, M., Witke, W., and Rottner, K. (2020). Molecular Dissection of Neurodevelopmental Disorder-Causing Mutations in CYFIP2. Cells 9.

62. Schneider, C.A., Rasband, W.S., and Eliceiri, K.W. (2012). NIH Image to ImageJ: 25 years of image analysis. Nat. Methods 9.

63. Singh, V., Davidson, A.C., Hume, P.J., Humphreys, D., and Koronakis, V. (2017). Arf GTPase interplay with Rho GTPases in regulation of the actin cytoskeleton. Small GTPases 0.

64. Singh, V., Davidson, A.C., Hume, P.J., Humphreys, D., and Koronakis, V. (2019). Arf GTPase interplay with Rho GTPases in regulation of the actin cytoskeleton. Small GTPases 10, 411–418.

65. Singh, V., Davidson, A.C., Hume, P.J., and Koronakis, V. (2020). Arf6 can trigger wave regulatory complex-dependent actin assembly independent of arno. Int. J. Mol. Sci. 21.

66. Steffen, A., Rottner, K., Ehinger, J., Innocenti, M., Scita, G., Wehland, J., and Stradal, T.E.B. (2004). Sra-1 and Nap1 link Rac to actin assembly driving lamellipodia formation. EMBO J. 23.

67. Su, L., Wang, Y., Wang, J., Mifune, Y., Morin, M.D., Jones, B.T., Moresco, E.M.Y., Boger, D.L., Beutler, B., and Zhang, H. (2019). Structural Basis of TLR2/TLR1 Activation by the Synthetic Agonist Diprovocim. J. Med. Chem. 62.

68. Sung, J.Y., Engmann, O., Teylan, M.A., Nairn, A.C., Greengard, P., and Kim, Y. (2008). WAVE1 controls neuronal activity-induced mitochondrial distribution in dendritic spines. Proc. Natl. Acad. Sci. U. S. A. 105.

69. Sztul, E., Chen, P.W., Casanova, J.E., Cherfils, J., Dacks, J.B., Lambright, D.G., Lee, F.J.S., Randazzo, P.A., Santy, L.C., Schürmann, A., et al. (2019). Arf GTPases and their GEFs and GAPS: Concepts and challenges. Mol. Biol. Cell 30.

70. Takenawa, T., and Suetsugu, S. (2007). The WASP-WAVE protein network: Connecting the membrane to the cytoskeleton. Nat. Rev. Mol. Cell Biol. 8, 37–48.

71. Tarricone, C., Xiao, B., Justin, N., Walker, P.A., Rittinger, K., Gamblin, S.J., and Smerdon, S.J. (2001). The structural basis of Arfaptin-mediated cross-talk between Rac and Arf signalling pathways. Nature 411.

72. Thompson, J.D., Higgins, D.G., and Gibson, T.J. (1994). CLUSTAL W: Improving the sensitivity of progressive multiple sequence alignment through sequence weighting, position-specific gap penalties and weight matrix choice. Nucleic Acids Res. 22.

73. Wang, J., and Hou, T. (2012). Develop and test a solvent accessible surface area-based model in conformational entropy calculations. J. Chem. Inf. Model. 52.

74. Wang, E., Sun, H., Wang, J., Wang, Z., Liu, H., Zhang, J.Z.H., and Hou, T. (2019). End-Point Binding Free Energy Calculation with MM/PBSA and MM/GBSA: Strategies and Applications in Drug Design. Chem. Rev. 119.

75. Wang, J., Wolf, R.M., Caldwell, J.W., Kollman, P.A., and Case, D.A. (2004). Development and testing of a general Amber force field. J. Comput. Chem. 25.

76. Wang, J., Hou, T., and Xu, X. (2006). Recent Advances in Free Energy Calculations with a Combination of Molecular Mechanics and Continuum Models. Curr. Comput. Aided-Drug Des. 2.

77. Wennerberg, K., Rossman, K.L., and Der, C.J. (2005). The Ras superfamily at a glance. J. Cell Sci. 118.

78. Yan, Y., Tao, H., He, J., and Huang, S.Y. (2020). The HDOCK server for integrated protein–protein docking. Nat. Protoc. 15.

79. Yin, J., Mobarec, J.C., Kolb, P., and Rosenbaum, D.M. (2015). Crystal structure of the human OX2 orexin receptor bound to the insomnia drug suvorexant. Nature 519.

80. Yin, J., Babaoglu, K., Brautigam, C.A., Clark, L., Shao, Z., Scheuermann, T.H., Harrell, C.M., Gotter, A.L., Roecker, A.J., Winrow, C.J., et al. (2016). Structure and ligand-binding mechanism of the human OX 1 and OX 2 orexin receptors. Nat. Struct. Mol. Biol. 23.

81. Zhang, Y., He, X., Zhai, J., Ji, B., Man, V.H., and Wang, J. (2021). In silico binding profile characterization of SARS-CoV-2 spike protein and its mutants bound to human ACE2 receptor. Brief. Bioinform. 22.

82. Zou, W., Dong, X., Broederdorf, T.R., Shen, A., Kramer, D.A., Shi, R., Liang, X., Miller, D.M., Xiang, Y.K., Yasuda, R., et al. (2018). A Dendritic Guidance Receptor Complex Brings Together Distinct Actin Regulators to Drive Efficient F-Actin Assembly and Branching. Dev. Cell 45, 362–375.e3.

83. Van Zundert, G.C.P., Rodrigues, J.P.G.L.M., Trellet, M., Schmitz, C., Kastritis, P.L., Karaca, E., Melquiond, A.S.J., Van Dijk, M., De Vries, S.J., and Bonvin, A.M.J.J. (2016). The HADDOCK2.2 Web Server: User-Friendly Integrative Modeling of Biomolecular Complexes. J. Mol. Biol. 428.

